# Substrate stiffness impacts early biofilm formation via a modulation of twitching motility

**DOI:** 10.1101/2022.02.18.480999

**Authors:** Sofia Gomez Ho, Lionel Bureau, Karin John, Delphine Débarre, Sigolène Lecuyer

## Abstract

Surface-associated lifestyles dominate in the bacterial world. Large multicellular assemblies, called biofilms, are essential to the survival of bacteria in harsh environments, and are closely linked to antibiotic resistance in pathogenic strains. Biofilms stem from surface colonization of various substrates encountered by bacteria. Here, we show that the opportunistic promiscuous pathogen *Pseudomonas Aeruginosa* explores substrates differently based on their rigidity, leading to striking variations in biofilm structure, surface decoration by exopolysaccharides (EPS), strain mixing during co-colonization and phenotypic expression. Using simple kinetic models, we show that these phenotypes arise through a mechanical interaction between the elasticity of the substrate and the type IV pilus (T4P) machinery, that mediates the surface-based motility called twitching. Together, our findings reveal a new role for substrate softness in the spatial organization of bacteria in complex microenvironments, with far-reaching consequences on efficient biofilm formation.

---

The transition of bacteria from a planktonic to a surface-attached state is of paramount importance in biofilm formation. In consequence, the way bacteria sense and respond to the close proximity of a surface has been the subject of intense scrutiny [1, 2]. This interaction involves different aspects of bacterial motility: swimming towards the surface, but also swarming, gliding or twitching that are used by attached bacteria to explore the surface collectively or individually [3, 4]. Eventually, permanent bacterial adhesion and colony structuration may arise, through mechanisms which essential ingredients are known (production of matrix, loss of motility), but in response to cues that remain unclear.

Bacteria are ubiquitous and can successfully colonize a wide range of biological tissues and abiotic surfaces [5, 6]. Different environments often result in different phenotypes for a given microorganism [7, 8]. However, although chemical signaling has long been known to impact bacterial gene regulation, it remains unclear how the mechanical properties of the encountered surface might impact bacterial behavior [9]. In this paper, we investigate how the rigidity of a substrate modifies bacterial motility, and by doing so impacts colony morphogenesis and early biofilm development. *Pseudomonas aeruginosa* (PA) is an opportunistic rod-shaped pathogen that contaminates a wide range of substrates, from very soft tissues to rigid implants [10, 11]. A particularly gifted and versatile biofilm-former, it is extremely prone to developing antibiotic resistance [12, 13]. PA has developed an arsenal of techniques to move on surfaces: among them is twitching motility, that allows single bacteria to translocate across surfaces using type 4 pili (T4P) [14]. T4P are thin protein filaments on the bacterial surface that can extend and contract by assembly and disassembly of the protein subunit PilA. The tip of T4P acts as a promiscuous hook that can grasp most surfaces. Attachment, contraction, detachment and extension cycles propel bacteria [14–17]. This surface motility is important for bacteria to efficiently settle on surfaces, but the exact mechanisms at play are unknown [18–20]. The function of T4P and the fact that it exerts forces on its environment make it an obvious candidate for surface-sensing mechanisms [15, 21, 22]. Recent results have shown that the polar localization of pili in PA could happen in response to surface-sensing [23]. Polarly-localized pili lead to persistent rather than random displacements, as well as specific effects such as the upstream migration of bacteria submitted to strong flows [24]. Reversal of twitching bacteria is rapidly induced upon meeting obstacles, suggesting a mechanical feedback from T4P [25]. In addition, PA can exert different types of virulence, from acute attacks to chronic infections [26, 27], and specific host-pathogen interactions have traditionally been considered as the key players in the regulation of these virulence pathways [28]. However, surface-sensing in itself has recently appeared as a potential signal that could trigger the upregulation of virulence-associated genes [29–31]. Although the global effect of surface rigidity on bacterial adhesion and biofilm formation has sometimes been adressed [32–34], so far how the micromechanical environment experienced by individual bacteria impacts their behavior is still unclear, possibly because of the difficulty to design and control microenvironments that allow for a fine tuning of mechanical properties at the bacterial scale, along with negligeable changes of the chemical environment.

In this study, we use a home-designed microfluidic setup to investigate at the single-cell level the influence of substrate rigidity on PA bacteria adhering to an open surface, under controlled flow conditions. We first investigate how substrate elasticity impacts early colony development. Focusing on single-cell behaviour, we study quantitatively how rigidity modulates bacterial motility and propose a purely mechanistic model to account for our observations. In turn, we demonstrate how this mechanical tuning of motility explains rigidity-induced changes in early surface colonization: colony morphology, matrix deposition, strain mixing and gene expression.

## RESULTS

In order to explore *in situ* the effect of substrate rigidity on the behavior of adhering bacteria, we have developed an experimental approach to include mechanically well-defined hydrogel pads in a microfluidic channel providing controlled flow conditions and allowing confocal imaging (Fig. 1A). We use the biocompatible hydrogel polyacrylamide (PAA), which has been extensively used to investigate cell-substrate interaction and mechanotransduction in mammalian cells. By varying the amount of bisacrylamide cross-linker during its preparation, PAA can span a biologically-relevant range of rigidities (from ∼ 1 to 100 kPa) while keeping a low viscous dissipation. Several pads of different rigidities, with Young’s modulus ranging from ∼ 3 to 100 kPa (see material and methods and Suppl. Fig. S12), were used in each experiment, and bacteria adhering on PAA pads and glass were imaged with high-resolution phase-contrast and fluorescence time-lapse imaging, from very low surface coverage up to the formation of microcolonies (1 frame/min over ∼ 10 hours).

**FIG. 1.**
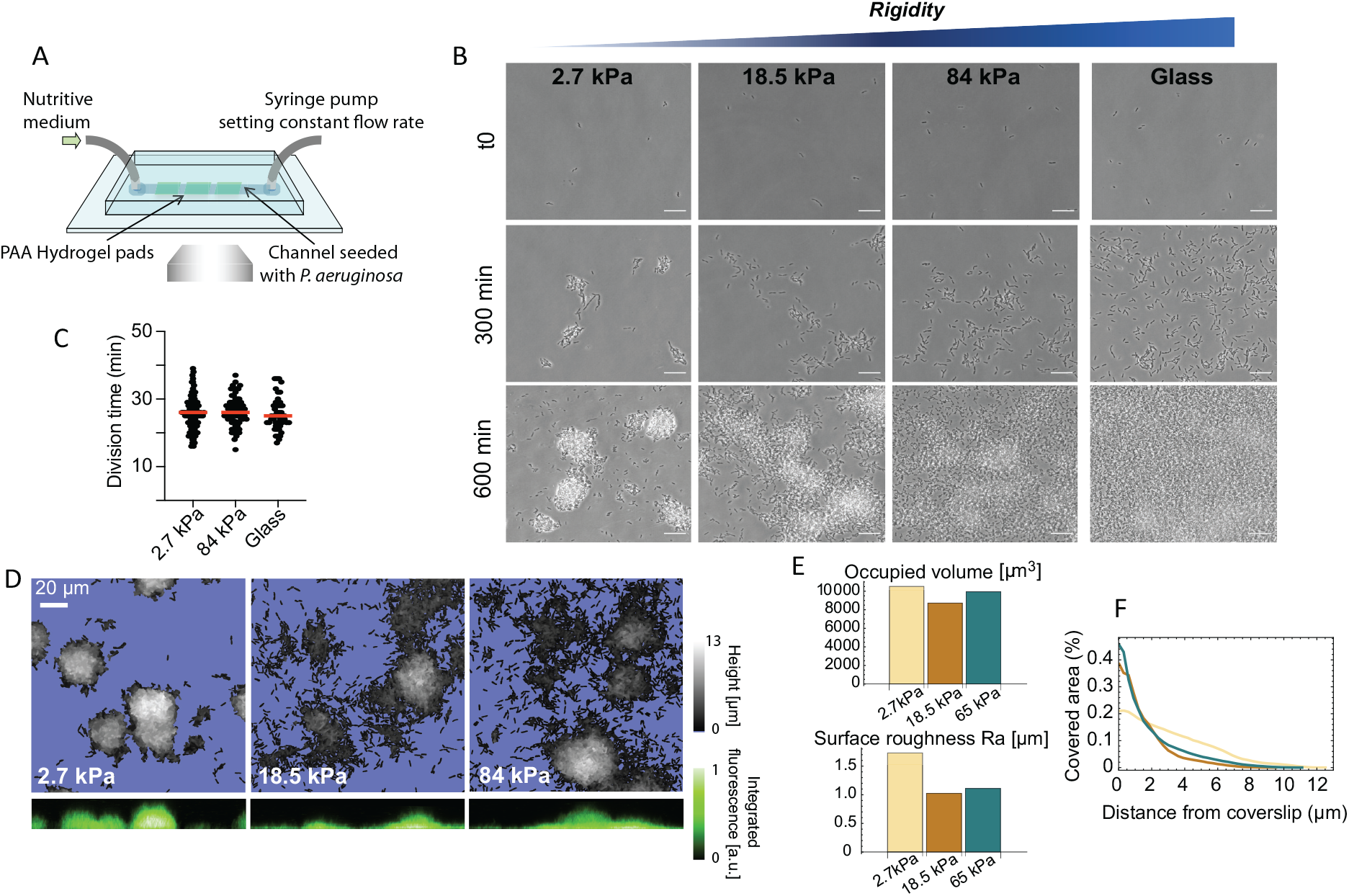
Bacterial microcolony formation depends on substrate rigidity. (A) Experimental setup: bacteria are imaged in a flow cell under constant flow of minimal medium. (B) After 10h, dense, isolated colonies form on soft PAA (2.7 kPa) while bacteria are more evenly distributed on stiff PAA, closer to what is observed on glass. Scale bars, 20 *µ*m. (C) Bacterial growth is not impacted by substrate rigidity. (D) 3D reconstruction of colonies confirms their hemispherical shape on soft substrates. (E) Total volume of colonies is conserved, but roughness is higher on soft substrates. (F) Fraction of area occupied by the bacteria as a function of the distance from the coverslip, showing flatter colonies on rigid substrates.

### Substrate elasticity modifies bacterial colonization of PAA in a T4P-dependent manner

We first focus on the effect of substrate rigidity on early colony formation. Straightforward observations with phase-contrast imaging show a striking impact on the shape of microcolonies after a few hours (Fig. 1B): on the softest hydrogels (¡ 10 kPa), bacteria form well-defined, dense hemi-spherical colonies; in contrast, on stiff hydrogels, bacteria are distributed in a thin layer covering most of the surface, a morphology closer to what we observe on glass. To rule out any effect of changes in the bacterial growth rate, we quantified the division time of bacteria (Fig. 1C), and the volume occupied by bacterial colonies after a few hours (Fig. 1D and E) on different substrates: both were found to be unaffected by the rigidity of the substrate, suggesting that bacteria develop and colonize substrates at the same rate irrespective of rigidity, but that the processes that drive their self-organization into colonies are modified. Indeed, in contrast, a change in the morphology of the colonies could be demonstrated by quantifying the characteristic roughness of the bacterial layer, which decreases as rigidity increases (Fig. 1E), and the distribution of bacteria with the distance from the surface, which spreads further for soft hydrogels (Fig. 1F). To control that this is a robust phenomenon driven by substrate elasticity rather than specific chemical interactions, we reproduced this assay using polyethylene glycol (PEG) hydrogels, which are chemically different from PAA but can span a similar range of rigidities. We obtained very similar results regarding the phenotype of colonies, which further confirms a role for the mechanical properties of the substrate in bacterial self-organization (Suppl. Fig. S6).

Next, we investigated how substrate rigidity could impact the morphology of early colonies: since surface motility is known to be important for initial self-organization of PA, we hypothesized that it could play a role in the different colony shapes that we observe. This link was explored by carrying out experiments using a mutant deprived of type IV pili (T4P), and thus unable to twitch on surfaces (mutant *PAO*1*pilA* :: *Tn*5, Suppl. Fig. S4). In these assays, the dependence of the colony morphology upon substrate rigidity is abolished and bacteria form dense hemispherical colonies on all PAA substrates. We therefore conclude that T4P-mediated surface motility (”twitching”) plays a key role in the rigidity modulation of microcolony formation of WT PAO1 on soft elastic substrates.

### Substrate elasticity modulates twitching motility efficiency

#### Experimental results -global motility

To quantify the coupling between the elasticity of the substrate and the twitching motility of bacteria, we analyzed time-lapse phase contrast images. These images allowed segmentation of individual bacteria (SI subsection I.A) from the starting time of the acquisition *t*_0_ (with a few isolated bacteria per field of view) until the transition to out-of-plane growth, after which bacteria cannot be easily separated anymore. From segmented binary images, we obtain the surface coverage *A*(*t*) as the fraction of occupied pixels, and the cumulative explored area *S* (*t*) as the fraction of pixels explored at any time up to time *t* (Fig. 2A).

**FIG. 2.**
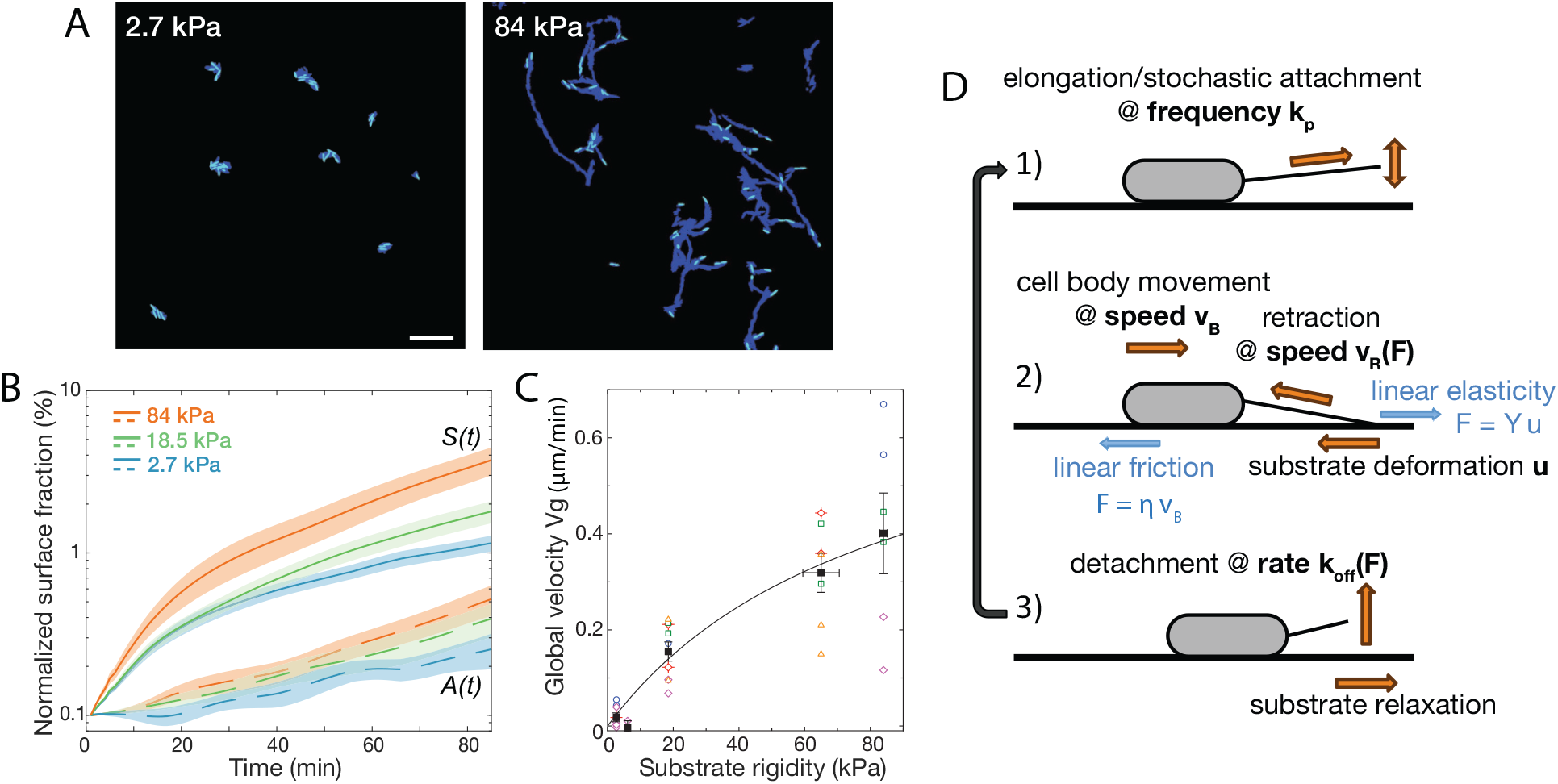
Bacterial surface motility is impaired on soft hydrogels. (A) Surface explored (dark blue) and current surface coverage (cyan) after 100 min on soft and stiff PAA surfaces. Scale bar: 20 *µ*m. (B) Surface coverage A(t) (broken lines) and cumulative explored area S(t) (full lines) for all tested rigidities (the initial surface coverage ¡A(t=0-10min)¿ was normalized to 0.1%). Shaded areas are standard errors of the mean (84 kPa: 6 data sets from 3 independent experiments, 18.5 kPa: 10 data sets from 5 independent experiments, 2.7 kPa: 8 data sets from 4 independent experiments).(C) Global bacterial motility *V*_*g*_ averaged over the first 100 min, inferred from the difference between A(t) and S(t) (16 different surfaces, 6 independent experiments). The black line is the fit with the kinetic model using equation 7. with values *V*_max_ = 0.77 ± 0.35*µ*m.min^−1^ and *E*_0_ = 84 ± 68 kPa. (D) Ingredients of the minimal 1D model for bacterial T4P-powered displacement.

The evolution of *A*(*t*) (Fig. 2B) reflects the exponential growth of initially attached bacteria on the surface, as well as potential attachment and detachment events during the acquisition. Ex-perimentally, because we are flushing the channel with clean medium, new adhesion events from free-swimming bacteria can be neglected at early times, so that

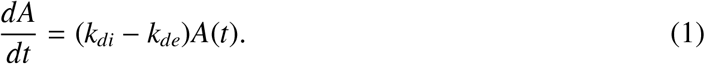

The bacterial division rate *k*_*di*_ does not depend on the substrate (Fig. 1C), and was measured for each experiment 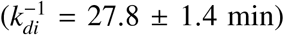. The difference observed in *A*(*t*) thus suggests that at short times, the detachment rate *k*_*de*_ is higher (although of the same order of magnitude) on softer hydrogels. Simultaneously, there is a quantitatively much larger dependence of *S* (*t*) on the substrate rigidity, which directly reflects the discrepancy in the effective motility *V*_*g*_ of bacteria on the surface. Indeed, the evolution of *S* can be written as

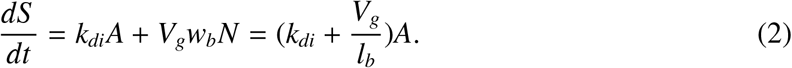

where *N* denotes the number of bacteria on the surface, and the typical size of a rod-shaped bacterium is *w*_*b*_*l*_*b*_ (width × length), so that the occupied area is *A* = *Nw*_*b*_*l*_*b*_. Here we have assumed that bacteria move along their major axis, neglecting reorientations (which is consistent with previous findings about polar localization of T4P [23, 25, 35] and our own observations).

The average bacterial length *l*_*b*_ was measured in each experiment (*l*_*b*_= 2.8 ± 0.13 *µm*). For each imaged position on a given substrate, we determined *dS* /*dt* and *A*(*t*) experimentally. The global bacterial velocity *V*_*g*_ was then estimated using Eq. 2, by averaging over the first 100 experimental time points.

During the first 100 minutes under flow, *V*_*g*_ exhibits a clear dependence on substrate elasticity (Fig. 2C and Suppl. Fig. S13A). Motility values are close to zero on very soft substrates (3-6 kPa), and progressively increase to reach ∼ 0.5 ± 0.25*µ*m/min on the stiffest hydrogels tested in this study (84 kPa).

#### Minimal kinetic modelling

A number of mechanisms could account for the experimental results presented above. Because the difference in twitching velocity is observed almost immediately upon attachment of bacteria onto the surface, we infer that phenotypic changes in response to surface-sensing might not be required to explain our observations. One simple hypothesis could thus be that a modulation of the twitching efficiency arises from the change in substrate elasticity through purely mechanical factors - the interplay between the T4P extension/retraction mechanism and the linear elasticity of the substrate - without the need for mechanotransduction mechanisms. To test this minimal hypothesis, we have developed a simple kinetic model, schematically described on figure 2D.

Briefly (more details can be found in SI section II), we consider a bacterium adhering onto an elastic substrate with a single effective pilus. The pilus is modelled as a rigid inextensible filament [36] and attaches to the substrate via its extremity with a typical adhesion size λ. The cell actively retracts its pilus until it detaches from the substrate with the force dependent velocity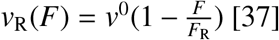, where *F* denotes the tensile load on the pilus, *F*_R_ the retraction stall force and *v*^0^ the retraction speed at zero load. Assuming linear elasticity, the tensile load *F* is related to the substrate displacement *u* at the pilus adhesion patch by *F* = *Yu*, where *Y* ∼ *E*λ and *E* is the Young’s modulus of the substrate. Since the typical size of the bacterial body *l*_*b*_ is much larger than λ, we neglect the deformation of the substrate induced by the bacterial body. Instead we assume that the pilus tension leads to a forward sliding of the bacterial body with a linear force-velocity relationship 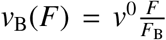(see SI subsection II.B and [38]), reducing the substrate deformation and the load in the pilus. Here, the ratio 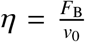 denotes the mobility constant of the cell on the substrate. With this model, the evolution of the pilus tension *F* is thus given by

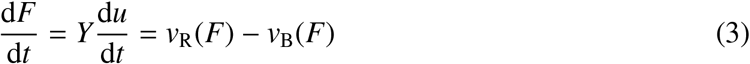

which is solved by

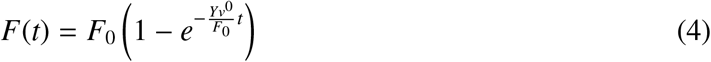

with the naturally arising force scale

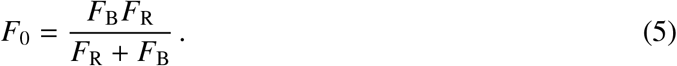

From <u>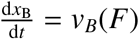</u>we obtain the bacterial sliding distance during the pilus retraction

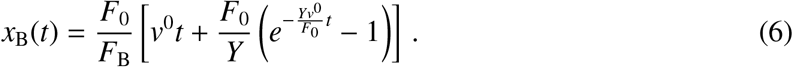

While retracting the pilus will detach with a rate constant *k*_off_ from the substrate. Assuming a force-independent off-rate constant 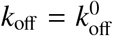(and hence a mean pilus adhesion time 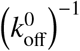 and a pilus retraction frequency *k*_p_, we obtain an effective bacteria velocity:

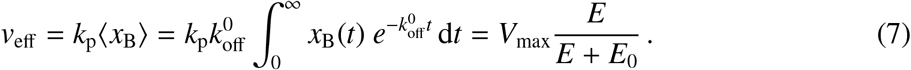

Here, ⟨*x*_B_⟩ denotes the mean bacterial sliding distance per pilus retraction event and *V*_max_ denotes the maximum effective speed a bacterium can reach on a given substrate at infinite rigidity. It is given by

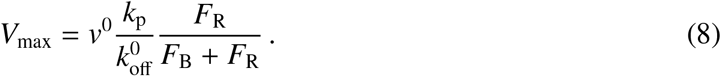

*E*_0_ denotes the rigidity at half-maximal speed and is given by

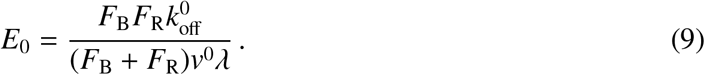

Fitting (7) against experimentally measured effective bacterial velocities *V*_*g*_ provides a quantitative description of the data for *V*_max_ = 0.77 ± 0.35*µ*m.min^−1^ and *E*_0_ = 84 ± 68 kPa. Conversely, we can estimate *V*_max_ and *E*_0_ from values of the parameters used in the model: assuming a typical pilus retraction speed *v*^0^ = 1 *µ*m.s^−1^ [37], a stall force of the order *F*_R_ = 100 pN [37], a pilus off-rate constant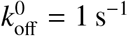[17], a contact size of λ = 1 nm, a high friction surface with *F*_B_ = 1 nN and a typical pilus retraction frequency[39] of *k*_p_ = 0.2 s^−1^ we obtain *V*_max_ ∼ 1 *µ*m.min^−1^ and a substrate rigidity at half maximum speed of *E*_0_ = 100 kPa, which are within 30% of the fitted values. Together this demonstrates that our experimental results on bacterial effective motility on elastic substrates can be interpreted as the result of a simple interplay between the pilus retraction mechanism, the deformation of the elastic substrate, and the friction of the bacterial body on this substrate.

#### Analysis of individual trajectories

The simple approach presented above to estimate an effective bacterial motility yields a population-averaged value of the velocity *V*_*g*_. Yet, the heterogeneity of bacterial populations has been highlighted in many studies, and is confirmed by our visual observations. Going further, we thus developed a segmentation and tracking protocol in order to measure the velocity of each individual cell at each time step of the acquisition (3A, see SI for details). This thorough approach yields the typical velocity distributions shown on figure 3A. These distributions show that bacterial displacements are indeed very heterogeneous, and can be well-fitted with an exponential decay so that the number of displacements with average velocity *V* is

**FIG. 3.**
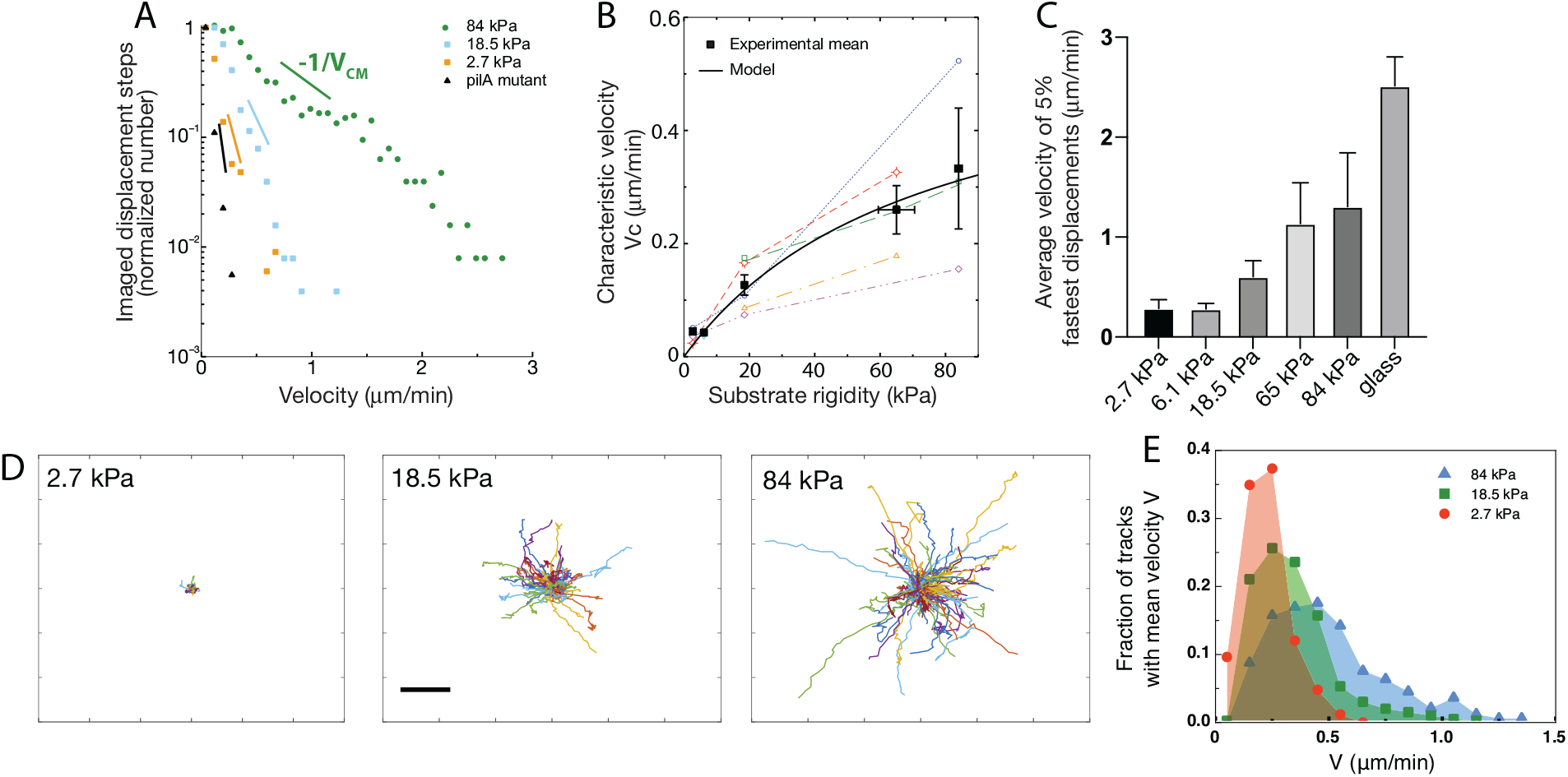
Twitching motility depends on substrate rigidity and is highly distributed in the bacterial population. (A) Normalized velocity distributions for the whole bacterial population on different PAA surfaces. The exponential decrease yields a characteristic center-of-mass velocity *V*_*CM*_ on each substrate. Displacement steps were measured every minute for 100 minutes, and two positions were acquired on each rigidity. The average of T4P-defective mutant on all surfaces is shown as a reference. (B) Active velocity characteristic values *V*_*c*_ obtained by fitting velocity distributions (6 independent experiments, 16 different surfaces). Values measured on different surfaces in a single experiment (same channel) are shown with the same symbols and connected. Black squares are mean values, and error bars show the SEM. The black line is the fit with the kinetic model using equation 7 with values *V*_max_ = 0.58 ± 0.31*µ*m.min^−1^ and *E*_0_ = 73 ± 72 kPa. (C) Average velocity values of the top 5% fastest imaged displacement steps for different substrates. Error bars are standard errors. (D) Individual bacterial tracks on soft (2.7 kPa), intermediate (18.5 kPa) and stiff (84 kPa) PAA during the first 3 hours after bacterial inoculation (total number of tracks is respectively 60, 123 and 175). Scale bar: 10 *µ*m (E) Mean track velocity distribution for different values of the substrate rigidity. Only full tracks were considered (corresponding to the right peak in Suppl. Fig. S7). Considering all tracks does not significantly modify the distributions (data not shown). 84 kPa: 330 tracks from 2 independent experiments, 18.5 kPa: 394 tracks from 3 independent experiments, 2.7 kPa: 83 tracks from 2 independent experiments.

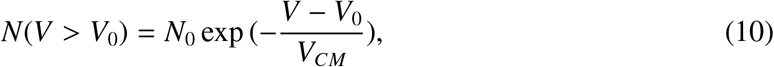

where *V*_*CM*_ is a characteristic velocity for the cell center of mass and *V*_0_ denotes a cutoff for low velocities (*V*_0_ = 0.08*µ*m/min). The pili knock-out strain was used as a reference for passive motility due to bacterial elongation, local reorganisations and experimental noise. This passive motility did not significantly depend on substrate rigidity (*V*_*CM*_(*pilA*) = 0.044*µ*m/min, see Suppl. Fig. S5), and its average value was subtracted from the values measured for the WT strain in order to selectively extract the characteristic velocity of active motility, *V*_*c*_. This analysis yields values for *V*_*c*_ (Fig. 3B and Suppl. Fig. S13B) in very good qualitative agreement with the global velocity analysis (*V*_*g*_, Fig. 2C). Again, our kinetic model describes the data quantitatively with values very close to the ones fitted and calculated in the previous subsection (*V*_max_ = 0.58 ± 0.31*µ*m.min^−1^ and *E*_0_ = 73 ± 72 kPa). In addition, we characterized the velocity of the 5 % fastest bacterial displacements (Fig. 3C). This analysis confirms the dependence of twitching velocity on substrate rigidity, but also yields higher velocity values, in good quantitative agreement with those reported in the literature using other experimental approaches [17].

Going further, we focused on the distribution of the measured displacements within bacterial tracks. Does a given bacterium alternate rapid and slow displacement phases, or do phenotypically distinct populations of slow and fast bacteria cohabit on the surface? To answer this question, we labeled each track, defined as the displacement of a cell between two division events (Fig. 3D). We first measured track duration, which we expected to be similar to the characteristic division time shown on Fig. 1C. However, we obtained a bimodal distribution with two peaks centered at times unaffected by the substrate rigidity: one peak is indeed centered on the division time (∼ 27 min), while the second one corresponds to bacteria spending 5 to 10 min on the surface before detaching (Suppl. Fig. S7). The velocity distribution corresponding to each peak is similar (data not shown). This observation is consistent with a phenotypical difference between daughter cells, in agreement with the results of [40] who showed that post-division, an asymmetric behaviour is observed with one daughter that keeps twitching while the other one leaves the surface. Interestingly, the fraction of bacteria that detach from the substrate is independent of the substrate rigidity (35 ± 2 %), but the duration of the track before detachment tends to be shorter on soft substrates. This duration is however difficult to interpret since it depends on the exact time of separation of the two daughter cells, which in turn depends on segmentation parameters that are arbitrarily defined in our protocol. We thus assume that after moving in sync with the first daughter cell, the second one often detaches from the substrate (about 70% of the time).

Focusing only on the adhering offsprings that remain on the surface, and normalizing tracks with respect to their initial position show a homogeneous radial distribution (Fig.3D), which confirms that shear does not influence bacterial orientation in our experiments. The distribution of the mean velocity of tracks does not allow us to distinguish different bacterial populations: it is broad, continuous with an exponential decay (reflecting the diversity of behaviours expected in a population of cells), and reaches higher values on stiffer hydrogels (Fig. 3E). In addition, for each track the standard deviation of this mean velocity is comparable and proportional to its mean (Suppl. Fig. S8), suggesting a stochastic distribution of the twitching steps within one trajectory.

### Rigidity-modulated bacterial motility governs the spatial characteristics of early surface colonization

*In-plane to 3D transition of emerging colonies*

To understand the way rigidity-modulated bacterial motility impacts the process of microcolony formation, we studied in details the way colonies transition to out-of-plane growth. Several experimental and theoretical approaches have been developed in the past to decipher this process: for confined colonies, the switch from planar to 3D growth takes place when it becomes energetically too costly to push neighboring cells outwards. In that case, the adhesion forces between the bacteria and their underlying substrate play a key role: strongly adhering bacteria transition to 3D colonies earlier in their development [41, 42]. In our experiments, there is no strong vertical or lateral confinement: bacteria can move on the substrate or away from it, so that cells stemming from a given progenitor do not necessarily stay in contact with each other. However, the twitching velocity determines how much cells, on average, move away from one another between two successive division events, thereby creating space to accommodate new offsprings on the surface.To investigate the possible link between twitching motility and 2D to 3D transition of growing microcolonies, we sought to determine *N*_*c*_, the number of adhered cells in a progeny (i.e. stemming from successive divisions of a given bacterium) when the 2D to 3D transition takes place. For softer substrates, all bacteria can be imaged, and *N*_*c*_ is directly measured; we also determined the average number of colonies per unit area. On stiffer substrates, it is impossible to track all bacteria stemming from a mother cell, since they are very motile and sometimes move out of the field of view. We assume that bacteria from other progenies are equally likely to move inside the field of view, so that measuring the number of bacteria on the image at *t*_*c*_, divided by the average colony density determined earlier gives a good approximation of *N*_*c*_. Fig. 4A shows *N*_*c*_ as a function of the center-of-mass characteristic velocity *V*_*CM*_ determined above (Fig. 3A), on different substrates and for 9 different experiments. *N*_*c*_ consistently increases with the twitching velocity, indicating a strong correlation between the twitching efficiency and the shape of early colonies and shedding light on our initial observations of variations in colony morphology as a function of the substrate rigidity (Fig. 1B, D-F).

**FIG. 4.**
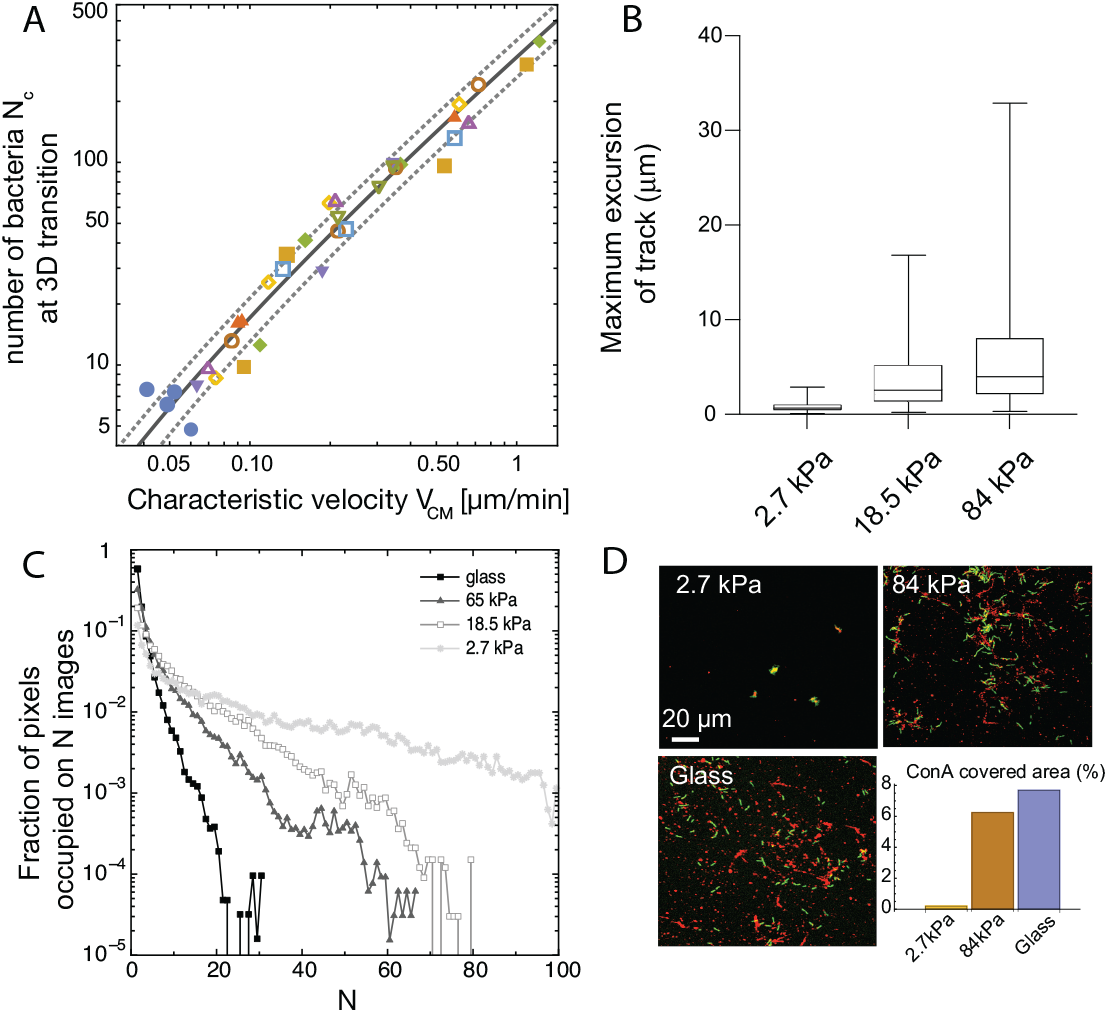
Spatial strucuring of surface colonization is impacted by substrate rigidity through twitching velocity. (A), size of microcolony (in number of bacteria *N*_*c*_) at the 2D to 3D transition as a function of the center-of-mass characteristic velocity *V*_*CM*_ defined in Fig.3A. Markers are experimental data from 9 different experiments, each with different substrates including glass (hence leading to higher values that in Fig. 3B). Blue dots are data obtained with the pili-deficient mutant pilA : Tn5. Lines, kinetic model for ⟨*γ*⟩ (solid line) ± its standard deviation (dotted lines). (B) Distribution of track lengths of full trajectories as a function of the substrate rigidity. (C) Distribution of occupation occurrence on each image pixel as a function of rigidity, showing a much more heterogeneous occupancy on soft subtrates. (D), ConA staining (red) of EPS deposition during cell (green) exploration of the surface.

To decipher the link between *N*_*c*_ and *V*_*CM*_, we have built a simple kinetic model with a single unknown parameter (SI section III). Briefly, we assume that the 2D to 3D transition takes place when the area occupied by bacteria reaches a fraction of the equivalent ”colony size”, defined as the characteristic area explored by bacteria in a progeny. Assuming that bacteria explore the surface through a random walk with persistence [37], the characteristic area accessible to bacteria in a colony over time can be written as *a*(*t*) = *a*_0_(1 + α*V*_*CM*_*t*) where *a*_0_ is the area of one bacterium and α is a parameter related to the properties of the random walk. Our experimental data show that not only the velocity, but also the contour length of the trajectories of bacteria increases with the rigidity since the duration of these trajectories are mostly constant (Fig. 4B and S7). Area *a*(*t*) is related, but not equal, to the area over which the colony spreads. Indeed, bacteria are not evenly distributed within the colony area, and we observe strong local density fluctuations. If we now consider an exponential growth of the number of bacteria on the surface due to the balance of cell division and detachment, it follows that the increase in the number of cells, and hence the area required to accommodate these cells on the surface, grows faster than the accessible area, driving a transition to 3D growth. Expressing the number of cells *N*_*c*_ in the colony at the time when this transition stochastically occurs leads to the following dependence as a function of *V*_*CM*_:

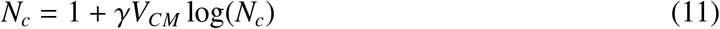

γ is an unknown parameter related to the properties of the random walk and the growth rate of bacteria on the surface that can be measured for each experiment. On figure 4A, we have plotted the corresponding curve using the average of experimental values for γ (solid line) ± their standard deviation (dotted lines). We observe an excellent agreement between this simple kinetic model and our experimental data over a wide range of velocities, including the T4P deficient mutant and the WT strain adhering on glass. This hints that it is the rigidity modulation of bacterial twitching velocity that shapes the organization of early colonies on elastic substrates, rather than energy minimization whereby bacteria would favour adhesion to the substrate or to other cells depending on the rigidity.

#### Surface decoration by extracellular matrix

One consequence of the modulation of twitching efficiency by the substrate elasticity could be the variation in matrix deposition by individual bacteria on the surface: indeed, upon adhesion, P. aeruginosa PAO1 secretes an extracellular matrix, mostly composed of exopolysaccharides (EPS), which was shown to result in the deposition of ”trails” on glass substrates. Since the matrix mediates the attachment of the cell body to the underlying substrate [43, 44], such deposits are inferred to facilitate further colonisation by bacteria and to impact colony formation. To investigate matrix deposition on hydrogel substrates, we introduced a fluorescent lectin (concanavalin A, see methods) in the nutrient medium infused in our device. The main component of PAO1 matrix, psl [45], is rich in mannose, that conA specifically binds [46]. Since conA interferes with the structure of the matrix, it was thus used for short-term imaging (t¡1h) of bacterial twitching at early stages, or added at the end of an acquisition (t∼ 8h) to assess matrix distribution on and around colonies. For high stiffness substrates, bacteria explore the surface efficiently, leaving trails of matrix and hence decorating a significant fraction of the surface; on the contrary, for nearly immobile bacteria on soft substrates matrix accumulates in confined areas, leaving most of the surface unmodified (Fig.4C and D). In addition, this difference in matrix distribution is maintained at a later stage of surface colonization (Suppl. Fig. S9). While on rigid hydrogels, most of the surface is covered by bacteria-secreted matrix, lectin staining on soft hydrogels is only present on compact colonies separated by regions completely devoid of EPS. While a number of possible experimental biases prevents interpreting the intensity of staining, quantification of the difference in EPS surface coverage highlights the impact of substrate rigidity on surface decoration by the bacteria.

### Substrate rigidity affects bacterial mixing

Real-life biofilms are generally formed by several species: pathogens can compete or help each other [**? ?** ], and commensal strains protect organisms from detrimental ones [47]. To further investigate how the variations in surface colonization with rigidity impact the structure of forming biofilms, we studied the model co-colonization of hydrogels by two PAO1 strains expressing different fluorescent proteins. Apart from their fluorescence expression, the two strains built on WT exhibit identical properties (motility, division rate, etc.). Through fluorescent confocal imaging, the strains can be spectrally separated to study their spatial distribution at different stages of surface colonization. As expected, rigidity-modulated motility impacts the co-colonization of the hydrogels from early stages (Suppl. Fig. S10): on rigid substrates, high motility promotes mixing of the offsprings of different cells, resulting in a spatial distribution of the two strains close to random (a residual correlation between the colour of neighbouring cells is always found due to the presence of cells that have just divided). Conversely, nearly immobile cells on soft substrates exhibit strong correlations between neighbouring cells which mostly arise from a single progenitor cell. This striking difference in strain mixing during surface co-colonization is maintained at later stages of biofilm formation: on soft substrates, quasi-monoclonal colonies with complete spatial segregation of green and yellow cells are observed, while the forming biofilms on rigid surfaces exhibit a close-to-random distribution of the two strains at the 10-*µ*m scale (Fig. 5A). To quantify this effect, we have used Moran’s I index, a statistical tool designed to quantify the spatial clustering of species. It provides a measure of the local spatial correlations and takes values ranging from 1 (perfectly correlated values) to -1 (perfectly anti-correlated values), with 0 corresponding to a spatially random distribution of the variable (see Supplementary Data I.C for details). The resulting quantitative analysis (Fig. 5B) confirms the decisive impact of rigidity on the structure of mixed biofilm with potentially far-reaching consequences on the interactions of different strains.

**FIG. 5.**
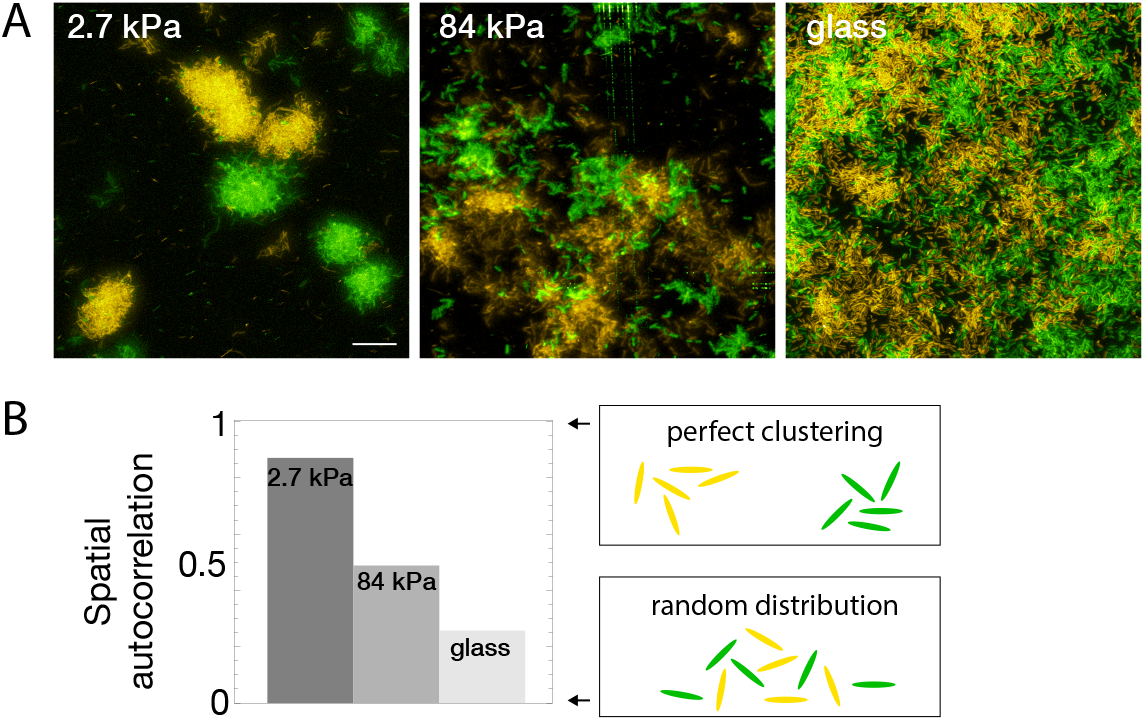
Bacterial spatial distribution is impacted by substrate rigidity. (A) Images of surfaces seeded with a 1:1 mixture of constitutively fluorescent bacteria expressing GFP or YFP show mostly monoclonal colonies on soft hydrogels, and mixed bacteria on rigid substrates (3D-rendering obtained by stacking images). Scale bar, 20 *µ*m.(B) Spatial correlations quantified via Moran’s I index.

### Surface rigidity impacts gene expression

Cell-cell communication, either via exported molecules or by direct contact is crucial during biofilm development [**?** ]. Modifications of bacterial distribution as described above thus likely impact gene regulation in surface-attached bacteria. To start addressing this complex question, we focused on the expression level of cyclic-di-GMP, a second messenger that controls the motile-to-sessile transition in P. aeruginosa [48]. We used the post-transcriptional fluorescent reporter build on the promoter of the gene *cdrA*, which encodes an exported protein involved in matrix cohesion, upregulated during biofilm formation by PAO1 [49]. The pcdrA-gfp intracellular reporter provides a measure of the integrated production of CdrA with a ∼ 40 min delay between expression of the gene and fluorescence detection [50]. Fig. 6 shows how *crdA* expression is modulated by the rigidity on 4 subtrates included in the same microfluidic device. For this reporter the degradation rate of GFP occurs over several hours, and its dilution due to growth and division of bacteria occurs at the same rate on all surfaces (see Fig. 1C). The increase rate of the fluorescent signal is thus a direct proxy to the expression rate of gene cdrA, and thus to the level of c-di-GMP.

**FIG. 6.**
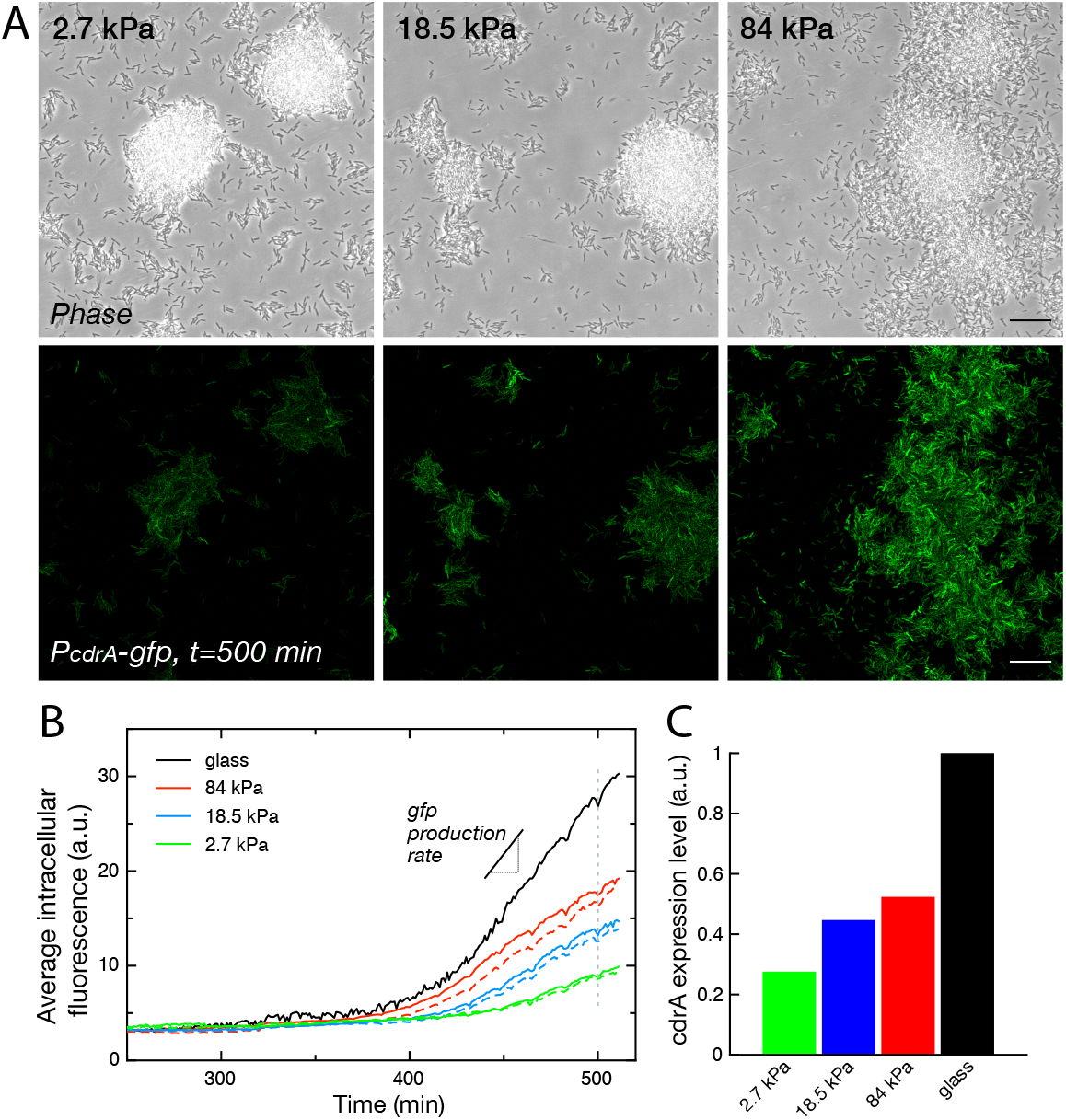
Substrate rigidity influences gene expression levels (A) Colonies grown from a modified PAO1 strain bearing the plasmidic *P*_*cdrA*_ − *g f p* reporter imaged in phase contrast (top) and fluorescence (bottom), after 500 min under flow. Scale bars, 20 *µ*m. (B) Average intracellular gfp fluorescence as a function of time, on different PAA surfaces and on glass. Broken lines are duplicate positions on a given surface. (C) Average cdrA expression level, obtained from a linear fit of (B).

During a first phase of surface colonization, fluorescence remains low on all surfaces. The signal subsequently starts increasing linearly, roughly at the same time for all surfaces (within the uncertainty of fluorescence quantification, i.e. ≈10 minutes). This second phase ends with the onset of a plateau, again around the same time for all surfaces, at the end of the exponential growth of bacteria adhered on the surface, possibly as a result of oxygen depletion in the flowing medium that would be sensed simultaneously on all surfaces (Suppl. Fig. S11). This linear increase in fluorescence directly translates into a constant production rate of CdrA that can be compared for the 4 surfaces (Fig. 6C): our analysis shows a marked increase in CdrA expression with the substrate rigidity.

## DISCUSSION

In this study, we have designed an experimental approach to investigate early microcolony formation by PAO1 on hydrogels with different elastic moduli, under constant flow rate. By continuously imaging surface-attached bacteria *in situ*, we show that substrate rigidity influences the twitching motility of individual bacteria, therefore strongly impacting the process of colony formation. Through two different analyses of the surface motility of the bacterial population, either via the global evolution of the explored area or via the tracking of individual cells, we find that the characteristic twitching velocity increases with substrate stiffness (from 0.02 to 0.7 *µ*m/min). The encounter between bacteria and a substrate generates interaction forces. Deciphering surface-sensing, i.e. understanding how these mechanical stresses generate signals that will translate into bacterial behavior has been the focus of a lot of recent research. It is now clear, for instance, that T4P contraction acts as a force sensor that transmits signals to the bacterium [51]. Here, however, our minimal mechanistic model suggests that a variety of observed phenomena (3D structure of colonies; EPS deposition on the surface; strain mixing during co-colonization) all derive from the modulation of the efficiency of pili activity by the deformability of soft substrates. We propose a 1D model based on a force balance between (i) a pilus that extends, attaches and retracts with a defined frequency; (ii) the deformation of the underlying substrate at the pilus tip upon retraction; and (iii) the friction force due to adhesion of the bacterial body when it is dragged across the surface at the other end of the pilus. In this balance, the detachment rate of the pilus tip from the substrate is a key parameter in the resulting bacterial velocity. When introducing the model we have assumed a force-independent off-rate constant for the pilus. In a more complex scenario, the contact between the pilus and substrate may act as a slip bond or a catch bond. For completeness we show some numerical results for slip and catch bond behavior in the SI (section I.D), which do not increase however the quality of fit between experimental and theoretical velocity data.

The strength and the originality of our model is that no mechanotransduction process is required: purely mechanical considerations are sufficient to account for our observations (twitching velocity, microcolony formation). Such mechanism may be of particular importance for surface colonization, since the adaptation of bacterial behaviour to its environment can thus be instantaneous. This process could complement mechanotransduction processes that might result in modulation of gene expressions - a process that require minutes to hours to take place (Fig. 6).

As an example of gene-modulation process, we have focused on c-di-GMP activation, via the gene *cdrA*: while the level of expression of the gene is clearly impacted by the substrate rigidity, differences in expression level are detected only 6-7 hours after the onset of surface colonization, with a first phase characterized by low *cdrA* expression on all surfaces. This timeframe suggests that the difference in gene expression that we observe is probably not due to a direct sensing of the substrate rigidity by individual bacteria, but rather a consequence of their organization into more or less dense colonies. Interestingly, our results do not show the classical association of high c-di-GMP level, increased matrix production and biofilm formation: matrix production and decoration of the surface are already present in the phase of low *cdrA* expression (Fig. 4D), and dense colonies also start forming during this phase on soft substrates (Fig. 1B). This hints that high c-di-GMP level is not necessarily required to trigger colony formation, that can instead be triggered by the micromechanical environment of the bacteria. In addition, while *cdrA* is thought to be upregulated in similar situations as matrix production (high c-di-GMP levels), here we observe an opposite trend (Fig. 6C and Suppl. Fig. S9): soft substrates seem to accumulate a high amount of EPS around dense colonies, while stiffer substrates are more extensively covered but with a lower local amount of EPS.

EPS distribution, composition and concentration may be significant for the recruitment of new cells on the surface: indeed, previously deposited matrix is thought to strengthen adhesion of *P*.*aeruginosa* bacteria [44], and could also possibly mediate adhesion of other microorganisms. A high amount of matrix in colonies growing on soft substrates likely increases the cohesion of the forming biofilm. Such process could reinforce the difference of surface colonization triggered by the rigidity-modulation of twitching efficiency: through a largely distributed matrix deposition, bacteria limit the increase in adhesion strength of newly-attaching bacteria to the decorated surface, and promote an evenly distributed density of cells on the surface. On the opposite, accumulation of matrix on dense colonies on soft substrates increases their cohesion and reduces further the possibility of outwards movement. This reinforcement process could be a way to resist detachment on a substrate less favorable to adhesion (see increased detachment rate at early stages of individual bacteria on soft substrates, Fig. 2B and Eq. 1).

In a wider context, this feedback loop could also be envisioned as an adaptive process to optimize bacterial colonization of mechanically heterogenous environments by ensuring accumulation of bacteria into dense colonies located in the softer regions of their environment, e.g. over cellular tissues. Recently, Cont et al. have shown that dense colonies were able to deform soft substrates and exert forces that could disrupt an epithelium layer [52]: rigidity-modulated twitching could thus provide *Pseudomonas Aeruginosa* with a convenient means of targeting soft tissues for cooperative disruption and subsequent invasion.

The phenotypic differences that we report are likely to impact subsequent interactions of bacteria with their environment: response to changes in nutrient or oxygen availability, and chemical signals in general which will not efficiently penetrate inside dense colonies. This could in particular impact susceptibility to antibiotics. This is all the more relevant that PA can invade many different environments, and might have to be treated differently when it settles in the lungs of cystic fibrosis patients, or on the surface of rigid implants.

Finally, our data show that rigidity-modulated twitching has a striking impact on the mixing of different strains upon surface colonization. Understanding the mechanisms governing the formation of mixed-species communities is one of the key challenges of current biofilm research. Since the motility modulation mechanism described here is quite general and should be marginally affected by the particulars of different strains/species moving through elongation/retraction of an appendage, we expect it to provide a relevant framework to study co-colonization in different mechanical micro-environments.

## MATERIAL AND METHODS

### Bacterial strains

Strains used in this study were Pseudomonas aeruginosa wild-type (WT) PAO1, fluorescent strains PAO1 mini-CTX-gfp and PAO1 mini-CTX-YFP, and PAO1 mutant pilA::Tn5 obtained from the transposon library at University of Washington [53]. Strain PAO1 pcdrA-gfp was obtained by transforming plasmid *pCdrA* :: *g f p*^*C*^ from [50] in our WT strain.

Bacteria were inoculated in Luria-Bertani (LB) medium from glycerol stocks, and grown overnight at 37 °C at 250 rpm. The next morning, 10 *µ*L of the stationary phase culture were diluted in 3 mL of LB medium and placed in a shaking incubator (37 °C, 250 rpm) for 3.5 hours, to reach midexponential phase (OD_600_ = 0.6-0.8). Bacteria were then diluted to OD_600_ = 0.005 in our working medium, TB:PBS, and inoculated into the channel. TB:PBS is obtained by mixing TB (Tryptone broth, Euromedex, 10 g.L^−1^) and PBS (w/o calcium and magnesium) with a ratio of 1:2. We found that this minimal medium favors bacterial twitching for a few hours after adhesion.

### Microfluidic device

Microfluidic channels were cut into 100 *µ*m-thick double-sided sticky tape (Teraoka, Japan) with a die-cutter. Typically, a 5 cm-long x 1 mm-wide channel was used to bind together a rectangular glass coverslip bearing hydrogel patches, and a flat 5 mm-thick slab of polydymethylsiloxane (PDMS, Sylgard prepared by mixing crosslinker and monomer solutions 1:10 and baking at 65°C for 1 hour). Two channels were stuck together to obtain a height of 200 *µ*m, in order for the flow through the channel to not be significantly modified by the 25 *µ*m-thick PAA hydrogels. The double sticky tape channels were first adhered onto the PDMS piece and then placed onto the dehydrated hydrogels. To insure proper binding, the whole device was placed under vacuum for 30 minutes. Next, the channel was rinsed with TB:PBS (1:2) for a minimum of 1 hr, in order to rehydrate the hydrogels. Medium was placed in a plastic container and withdrawn into the channel with a syringe pump (Harvard Apparatus, USA, 30 *µ*L/min) to avoid the formation of bubbles.

### Gels and substrates preparation

Hydrogels of polyacrylamide (PAA) and polyethyleneglycol (PEG) were prepared following previously established protocols [54, 55]. All reagents were obtained from Sigma Aldrich and used as received: Acrylamide solution (AA, 40% in water), N,N’-Methylenebisacrylamide (Bis, 2% in water), Ammonium Persulfate (APS, ≥ 98%), N,N,N’,N’-Tetramethylethylenediamine (TEMED, ≥ 99%), Poly(ethylene glycol) diacrylate (PEGDA, *M*_*n*_ ∼ 6000 g.mol^−1^), 2-Hydroxy-4’-(2-hydroxyethoxy)-2-methylpropiophenone (Irgacure 2959, 98%), Bind-silane, Sigmacote.

Rectangular glass coverslips (24×60 mm) were used as substrates for gel casting. They were plasma-cleaned and immersed in a solution of Bind-silane (60 *µ*L of Bind-silane, 500 *µ*L of 10 % acetic acid, 14.5 mL of ethanol) for 1 hour before being rinsed with ethanol and water, and blow-dried with nitrogen before use. Round glass coverslips (12 mm diameter) were used as counter-surfaces for gel casting. After plasma cleaning, they were immersed in Sigmacote for 1 hour before rinsing with acetone, ethanol and water, and blow-dried before use.

Bulk solutions of AA/Bis and PEGDA were prepared in phosphate buffer saline (PBS) and stored at 4 °C until use. The final stiffness of the gels was tuned by adjusting the AA/Bis or PEGDA content according to Table I. PAA gels were obtained by adding 1 *µ*L of TEMED and 1 *µ*L of a freshly made APS solution (10 w% in water) to a volume of 168 *µ*L of AA/Bis solution. A 3 *µ*L droplet of the mixture was immediately placed on the surface of a bindsilane-treated glass coverslip, sandwiched by a Sigmacote-treated round coverslip, and left for curing for 1 hour in a water vapor-saturated atmosphere. After curing, the round coverslip was lifted off using the tip of a scalpel blade, resulting in a circular pad of gel, of thickness ∼ 25 − 30 *µ*m, covalently bound to the bottom rectangular coverslip and exposing its free top surface. Circular gel pads were then scrapped with a razor blade in order to adjust their lateral size to the width of the microfluidic channels into which they would eventually be installed. Gel pads were then copiously rinsed with ultrapure water, and left for drying in a laminar flow cabinet. Up to two such pads, with different elastic properties, were prepared simultaneously on the same coverslip, arranged to fit along the length of the microfluidic channels. PEG gels were obtained by adding 5 *µ*L of a 10 wt% solution of Irgacure in ethanol to 0.5 mL of PEGDA solution. A 3 *µ*L droplet of the mixture was placed in between coverslips as described above, and irradiated under UV light (365 nm, 180 mW.cm^−2^) for 15 minutes for curing. Subsequent steps were as described above for PAA gels.

**TABLE I.** Hydrogel compositions and associated Young’s moduli

| Acrylamide (wt%) | Bis-acryl. (wt%) | PEGDA (wt%) | Modulus (kPa) |
| --- | --- | --- | --- |
| 4 | 0.225 | 0.0 | 2.7±0.3 |
| 5 | 0.225 | 0.0 | 6.1±0.2 |
| 8 | 0.264 | 0.0 | 18.5±0.7 |
| 20 | 0.47 | 0.0 | 65±5.6 |
| 15 | 0.65 | 0.0 | 84±1.1 |
| 20 | 0.7 | 0.0 | 103±3.8 |
| 0 | 0 | 5 | 5.7±0.3 |
| 0 | 0 | 20 | 102±8.4 |

### Mechanical characterization

The viscoelastic properties of the various gels were characterized by AFM microrheology, using the “contact force modulation” technique described recently and validated on hydrogels [56]. It allows determining elastic and loss shear moduli, *G*′ and *G*″, as a function of frequency over the range 1 − 300 Hz. The Young moduli reported in table I have been computed as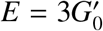, with 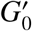the low frequency plateau modulus obtained by microrheology, assuming a Poisson ratio *v* = 0.5 for all gels. All gel samples displayed elastic behavior with *G*′ ≫ *G*″.

Measurements were performed on a JPK Nanowizzard II AFM, with pyramidal-tipped MLCT probes (Bruker) of spring constant 15 mN/m. Data were analyzed using a home-written software for microrheology. 30*µ*m-thick gels were prepared, as described above, on round coverslips mounted at the bottom of 35 mm petri dishes. They were then either characterized immediately or left to dry to mimick the protocole used for inclusion in the flow chamber. Experiments were performed in PBS + 1 % vol. tween 20 (Sigma), with Tween used to prevent adhesion of the AFM tip to the gel. All measurements were carried out at 37 °C to mimick experimental conditions with bacteria. Results were compared with force-distance indentation curves that gave consistent results at low rigidities (< 20 kPa) but overestimated the rigidity for higher values (see Suppl. Fig. S12).

Homogeneity of the gels was assessed at the *µ*m and mm scales by multiposition measurements. We found very good repeatability of the measurements and homogeneity of the gels at all scales (Suppl. Fig. S12). Subsequent measurements were hence acquired at 3-6 different points in the gels and the average and standard error of the mean are provided (Table I). Rigidity was also measured before and after drying and rehydration of the gel to check for possible damage to the structure. In addition, confocal images of the surface of fluorescently labelled gels were used to track default on the gel surface before and after drying. We found no evidence of damage to the hydrogel upon drying, except for very soft gels of rigidity below 1 kPa that were not used in this study (Suppl. Fig. S12).

### Microscopy experiments

Diluted bacterial solution was pumped into the channel, and kept without flow for 30 min to allow bacteria to attach. During that time, inlet tubing was connected to a syringe filled with TB-PBS medium supplemented with 3 mM glucose. 30 min after injecting bacteria into the device, the flow of medium was initiated. The flow rate was first set at 20-30 *µ*l/min for 2-3 min in order to flush out unattached bacteria, and then lowered to 1 *µ*L/min and maintained constant with a syringe pump (Pico Plus, Harvard Apparatus) throughout the acquisition. The set up was placed into the incubation chamber (37^*o*^C) of a Leica SP8 confocal microscope, and acquisition was started at 1 frame/minute.

## Supporting information

supplementary informations

## ACKNOWLEDGMENTS

We thank Claude Verdier for help with the AFM elasticity measurements, Benoit Coasne and Benedikt Sabass for fruitful discussion on data modeling, More thanks should go here. D.D. was supported by the French National Research Agency (grant ANR-19-CE42-0010). The authors acknowledge support from LabeX Tec 21 (ANR-11-LABX-0030).

