## supplementary informations for "Substrate stiffness impacts early biofilm formation via a modulation of twitching motility"

### Supplementary Information

#### I. IMAGE ANALYSIS

All image processing and analysis, unless otherwise noted, was performed with Fiji using available plugins and home-written macros.

##### A. Tracking of individual bacteria

In order to quantify the movements of individual bacteria, time series of phase-contrast images were registered using the plugin "MultiStackReg" [59] and segmented with the plugin "weka trainable segmentation" [60]. The resulting segmentation was then checked and corrected manually. The "analyze particle" function was subsequently used to locate the center of mass of each bacteria. Finally, the plugin "TrackMate" [61] was used to track all individual bacteria, again followed by manual validation and correction.

##### B. Quantification of the morphology of colonies

Quantification was performed on confocal fluorescence 3D resolved images. First, signal attenuation with depth was compensated by decreasing exponential fitting of the mean pixel values inside the colony with depth, and normalization by the corresponding function. A 2D 3x3 smoothing operation was then performed on each image of the z-stack, and the colonies were subsequently segmented using a simple thresholding operation: while this procedure does not permit segmentation of individual bacteria, it provides a good estimate of the 3D envelope of the colonies. The topology of the colonies was then quantified by calculating the roughness of this envelope using the widely-used arithmetic average roughness $Ra$

$$Ra = \frac{1}{N}\sum_{i=1}^{N}|z_i - \langle z\rangle| \, , \tag{1}$$

where summation is over all 2D positions $i$ in the 3D image, $z_i$ is the height of the highest segmented pixel at position $i$ and $\langle\rangle$ is the averaging operator over all positions. The occupied volume $V$ is calculated as

$$V = px^2 \sum_{i=1}^{N} z_i \, , \tag{2}$$

with $px$ the pixel size. The occupied area as a function from the distance to the coverslip is the histogram of $z_i$ values with bin size 0.5 µm (corresponding to the vertical sampling of the 3D images).

##### C. Quantification of the mixing of two strains co-colonizing the same soft substrate, as a function of the softness

This quantification is performed both at the low density stage with isolated bacteria, and at a later stage on maturing colonies. To this aim, we used a statistical tool, Moran's I index, designed to quantify the spatial clustering of species and widely used in the field of ecology and geography [62]. Moran's I is a measure of the local spatial correlations that includes a notion of spatial proximity, either in the form of a spatial cut-off for the calculation of the heterogeneity (in other words, a characteristic distance), or a number of neighbors. It takes values ranging from 1 (perfectly correlated values) to -1 (perfectly anti-correlated values), with 0 corresponding to a spatially random distribution of the variable.

Considering a variable $y$ that can take two different values (in our case, green (1) or yellow (-1)) with $n$ realisations, Moran'I is expressed as:

$$I = \frac{n}{\sum_{i=1}^{n}\sum_{j=1}^{n} w_{ij}} \frac{\sum_{i=1}^{n}\sum_{j=1}^{n} w_{ij}\left(y_i - \langle y\rangle\right)\left(y_j - \langle y\rangle\right)}{\sum_{i=1}^{n}\left(y_i - \langle y\rangle\right)^2}, \tag{3}$$

where $w_{ij}$ is the matrix of weights that contains the spatial information (with $w_{ii} = 0$). In our experiment, the relevant spatial scale (and hence the matrix of weights) varies greatly over time because of the change in the density of the bacteria on the surface. While at high density (maturing colonies) defining a length scale is a suitable way of testing the presence of local correlations, this is more challenging at earlier times when the distance between neighbours exhibits large stochastic variations, in particular for stiff substrates. Hence, different matrices of weights were chosen for early-stage and later-stage colonisation of the surface:

- at early stages of colonisation, when the bacteria are sparse on the surface, we chose to focus on the nearest neighbours of each bacteria. To this aim, individual bacteria are segmented in the green and yellow images, and their center of mass location is collated into a list of 2D coordinates and colour for all bacteria in the field of view. Moran's I is then calculated based on this list using the following weight matrix:

$$w_{ij} = \begin{cases} 1 & \text{if } j \text{ is one of the } p \text{ nearest neighbours of bacteria } i \\ 0 & \text{otherwise} \end{cases}$$

  We arbitrarily chose $p = 5$ as a significant number of neighbours, although similar results are found for values of $p$ ranging from 4 to 10. Lower numbers are biased by cell division: at the time of division, the closest neighbour is necessarily of the same strain as the bacteria under consideration, so that there is always a positive correlation between them. As a result, testing for mixing requires to mitigate this effect by choosing a large enough value for $p$. In practice, we found that $p = 5$ was a good compromise to limit this bias while maintaining a "local" approach, i.e. not considering the correlation between bacteria further apart than half of the field of view (i.e. 160 µm).

- at later stages with dense, 3D colonies, individual segmentation of bacteria becomes challenging and the correlation measure is performed on individual pixels: first, a simple thresholding operation is performed on the green and yellow image, and each pixel is attributed a value: 1 (green pixel), 0 (black pixel) or -1 (yellow pixel). From this new image, Moran's I is calculated using the following weight matrix:

$$w_{ij} = \begin{cases} 1 & \text{if the distance between } i \text{ and } j\,(i \neq j) \text{ is smaller than or equal to } d \\ 0 & \text{otherwise} \end{cases}$$

  Again, the cut-off distance $d$ is arbitrarily chosen as 5 µm although values between 3 and 10 µm yield similar results: it permits limiting fluctuations by averaging over a significant number of bacteria, while maintaining a local measure of mixing. In addition, because individual bacteria cover more than one pixel in the acquired images, a number of pixels of the same colour as pixel $i$ are removed to avoid correlating the bacteria with itself. In our data the average number of pixels covered by one bacteria is measured to be 40.

While there is some degree of freedom on the choice of the weight matrix, it is important to note that we use the same weight to compare data obtained on three different rigidities, hence minimising the impact of the exact chosen parameters on the comparison. In contrast, values obtained on one surface at the two different time points should not be directly compared as they have not been obtained with the same weight matrix.

#### II. MODELING TWITCHING VELOCITIES ON SOFT SUBSTRATES

The principle of our modeling of rigidity-modulated twitching in 1D is shown in figure 2D (main text), and incorporates three main ingredients: modeling of the substrate deformation (subsection II A), of cell body friction on the surface (subsection II B) and of the pilus retraction dynamics (subsections II C and II D).

##### A. Modeling substrate deformation

We have based our approach on the theory of linear elasticity for the description of the substrate: in this framework, the deformation of the substrate occurs over a typical length scale given by the size of the adhesion, $\lambda$, and it is proportional to the force applied. Finally, the proportionality coefficient $Y$ scales as the product of the substrate elastic modulus, $E$, and the adhesion size $\lambda$, i.e. $Y = E\lambda$. This simple relation is valid only for small displacements on rigid substrates. It is likely to fail quantitatively on very soft substrates with large displacements, low cross-linker densities and non-affine deformations, but is a reasonable first approximation for the simple model we propose here.

This modeling introduces characteristic length scales that depend on the part of the bacteria under consideration: both the pilus and the cell body form contact with the substrate. The pilus attaches at its tip over size $\lambda \approx 1$ nm, while the cell body has a typical size of $l_b \approx 1\mu$m. In addition, a third length scale is the typical length of the pilus, $L$, which varies during retraction but is most of the time $> 1\mu$m. Introducing these three quantities permits to simplify the description of the deformation of the substrate: the pilus tension $F$ and the displacement at the adhesion site in the substrate $u$ are linearly related by $F = Yu$, with $Y$ being an effective spring constant. We model the substrate as an infinite (thickness$\approx 25\,\mu$m$\gg \lambda$, lateral extension$\approx 1 - 10\,$mm$\gg \lambda$), isotropic, elastic and incompressible half space. Furthermore, we neglect the influence of the cell body on the deformation around the pilus tip since $L >> \lambda$ so that the deformation of the substrate has decayed to zero at the cell body.

The 2D Boussinesq Green's tensor at the surface $z = 0$ for a point like shear force $\mathbf{f}$ at the origin is given by [63]

$$\mathbf{G}(\mathbf{r}) = \frac{3}{4\pi E}\left[\frac{\mathbf{I}}{r} + \frac{\mathbf{r}\otimes\mathbf{r}}{r^3}\right]\mathbf{f}\,. \tag{4}$$

Considering an adhesive type IV pilus (T4P) tip of length $\lambda$ and half-width $d$ and using slender body approximations, the total force $F$ on the pilus for a "lengthside" displacement $u$ is given by

$$F = \frac{E\lambda\pi}{3\ln\frac{\lambda}{d}}u \quad \text{with} \quad Y = \frac{E\lambda\pi}{3\ln\frac{\lambda}{d}} \approx E\lambda\,. \tag{5}$$

Here we have implicitly introduced a 1D setting, i.e. we will neglect the vectorial nature of forces and displacements and restrict ourselves to a 1D setting. We find, as expected, that $Y$ scales linearly with $\lambda$. This holds equivalently for the cell body by replacing $\lambda$ with $l_b >> \lambda$: as a result, the substrate deformation at the cell body caused by the same pilus tension $F$ is of amplitude smaller by a factor $\lambda/l_b \ll 1$ and will be neglected for the sake of simplicity.

In contrast, we consider the pilus tip to be firmly attached to the substrate until detachment while the cell body can slide on the surface. Note that this asymmetry between bacterial body (macroscopic sliding over the substrate) and the supposedly small pilus/substrate contact (point-like force deforming the substrate) is the essential difference to the pulling process described in Ref. [64], where the contact of the bacterial body and the pilus extremity are mechanically treated as equivalent.

##### B. Modeling cell body friction

As stated above, the model requires a description of the sliding motion of the cell body on the substrate as a function of the force $F$ applied by the pilus. We base our modeling on the theory from [65] that considers stochastic friction by an ensemble of N elastic linkers (not necessarily all bound at all times) between an elastic substrate and a cell, submitted to a sliding velocity $v$. The bonds are modeled as slip bonds with a critical force $f^*$, an off-rate constant at zero force $k^0_{\text{off}}$ and an on-rate constant $k_{\text{on}}$. The linkers' stiffness is $k_{\text{b}}$.

In the case of an infinitely rigid substrate, the mean total force on the cell body $\langle F\rangle$ as a function of its velocity $v$ is non-monotonous and is given by

$$\langle F\rangle = Nf^* \frac{r_{\text{on}}\, e^{1/\tilde{v}} \int_0^\infty f e^{-\left(\frac{e^f}{\tilde{v}}\right)} \mathrm{d}f}{\tilde{v} + r_{\text{on}}\, e^{\left(\frac{1}{\tilde{v}}\right)}\Gamma\left[0, \frac{1}{\tilde{v}}\right]} \tag{6}$$

with $\tilde{v} = v/v_\beta$, $v_\beta = k^0_{\text{off}} f^*/k_{\text{b}}$, $r_{\text{on}} = k_{\text{on}}/k^0_{\text{off}}$. $\Gamma\,[0, x]$ is the Euler gamma function. Eq. (6) exhibits a complex dependence of $\langle F\rangle$ on $\tilde{v}$ that requires estimating typical values of the parameters in our experiments. Putting in numbers to obtain the typical speed $v_\beta$, we can estimate that

- $k^0_{off} \approx 1 - 10\ s^{-1}$ (slightly higher than for specific ligand/receptor bonds [66])
- $f^* = k_{\text{B}}T/x_\beta$ with $x_\beta \approx 0.1 - 10\,$nm being the transition state distance between bound and unbound state as proposed by Evans [67] and others [68].

- $k_b$ is more difficult to estimate. Here we assume that bacterial adhesion is mediated by the bacteria produced extracellular matrix, of which a major constituent are exopolysaccharides. Using a worm-like chain (WLC) model for a polymer of persistence length $L_\mathrm{p} \approx 10\,\mathrm{nm}$ (as calculated for bacteria produced exopolysaccharides in [69]) and contour length $L_0 \approx 100\,\mathrm{nm}$ (assuming a chain length of about 100 monomers with size 1 nm), the linear force-elongation relationship in the regime of weak forces [70] yields a force constant $k_b \approx \frac{3kT}{2L_\mathrm{p}L_0} \approx 6\times10^{-3}\,\mathrm{pN.nm^{-1}}$.

Taking extreme values this leads to typical velocities in the range $v_\beta = 1 - 100\,\mathrm{\mu m.s^{-1}}$. In our experiments the bacteria are not expected to move faster than the pilus retraction velocity (i.e. $1\,\mathrm{\mu m.s^{-1}}$ [71], if one excepts the case of slingshots that were not frequently observed in our experiments). Taking into account that the pilus retraction speed slows down considerably as the tension in the pilus increases, the bacterial speed during one pilus retraction is rather smaller than this maximum value. Hence, we always have $\tilde{v} = v/v_\beta < 1$, and Eq. (6) can be linearized to

$$\langle F\rangle = Nf^* \frac{k_\mathrm{on}}{k^0_\mathrm{off} + k_\mathrm{on}}\tilde{v}, \tag{7}$$

In addition, the elasticity of the substrate should be considered. Ref. [65] proposes that this situation is equivalent to having a system of springs in series, one stemming from the substrate elasticity and the second being the collection of individual bond springs (in parallel). In this case and using once again the theory of linear elasticity, the previous analysis holds if $\tilde{v}$ is rescaled by a factor $\frac{E\,l_b}{k_b+E\,l_b}$, with $E > 3\,\mathrm{kPa}$ the substrate Young's modulus and $l_b \approx 1\,\mathrm{\mu m}$ the characteristic size of the bacterial cell body, $a = l_b \approx 1\,\mathrm{\mu m}$. Hence $El_b \geq 3\,\mathrm{pN.nm^{-1}} \gg k_\mathrm{b}$ and the scaling factor $\frac{E\,l_b}{k_\mathrm{b}+E\,l_b} \approx 1$, so that the elasticity of the substrate does not influence the friction of the cell body.

In summary, we find that we can reasonably use a linear approximation for the bacterial sliding speed in response to the pulling force due to the pilus retraction, $F = \eta v$ with $\eta$ a friction coefficient. Finally, we consider $\eta$ as independent from the substrate rigidity, which is reasonable if we assume that the number of bonds is limited by the number of molecules/appendages of the cell body that can interact with the substrate, rather than the number of binding sites on the substrate itself (PAA mesh size $\approx 3 - 10$nm), and that the interaction may in addition be mediated by adsorbed exopolysaccharides deposited by the bacteria. However, other non-linear dependencies can be easily included into the modeling.

##### C. Basic modeling of pilus retraction

The relevant step during twitching which induces bacterial motion is the active pilus retraction when attached to the substrate. Here we assume, that the limiting effect for bacterial motion is the detachment of the pilus from the substrate, and not the complete retraction of the pilus by the bacterium. To understand the role in substrate rigidity on the bacterial twitching speed we will therefore concentrate on this crucial step without describing the whole cycle of pilus dynamics, for which the kinetics is not completely understood [72, 73].

We consider the retraction of a single effective pilus pulling on the bacterial body until it detaches from the substrate. We treat the pilus as rigid and inextensible filament: assuming a force constant of $2\,\mathrm{pN.\mu m^{-1}}$ for the pilus elasticity [74], a substrate rigidity of $E = 100\,\mathrm{kPa}$, an adhesion size of $\lambda = 1\,\mathrm{nm}$ and a maximum force exerted by the pilus of $F_\mathrm{R} = 100\,\mathrm{pN}$, the substrate displacement is $u \sim F_\mathrm{R}/E\lambda = 1\,\mathrm{\mu m}$. In contrast, the pilus elongation is $\Delta L = 50\,\mathrm{nm}$ and can therefore be neglected for our conditions. However it would not pose any difficulty to include the pilus elasticity into the calculations.

Let $v_R$ be the retraction speed of the attached pilus inducing a displacement $u$ in the substrate. At the same time the bacterium will slide forward with speed $v_B$, reducing the tension in the pilus and the displacement in the substrate:

$$\frac{\mathrm{d}u}{\mathrm{d}t} = v_\mathrm{R}(F) - v_\mathrm{B}(F) \quad \text{with} \quad F = Yu\,. \tag{8}$$

Both motions (substrate displacement and bacterial sliding) are coupled via the tension in the pilus $F$. Its retraction speed is described by a simple linear dependence that has been well documented [75, 76]

$$v_\mathrm{R} = v^0\left(1 - \frac{F}{F_\mathrm{R}}\right), \tag{9}$$

with $F_R$ a stall force. As established in the previous subsection, the bacterial sliding speed depends linearly on the pilus tension with friction constant $\eta = F_B/v^0$:

$$v_B = \frac{1}{\eta} F = v^0 \frac{F}{F_B} \,. \tag{10}$$

$F_B$ denotes the force necessary to pull the bacterium at maximum retraction speed $v^0$ over the substrate. From Eq. 8 we recover the increase in the pilus tension over time during the retraction

$$F(t) = F_0 \left(1 - e^{-\frac{Y v^0}{F_0} t}\right) \tag{11}$$

with the force scale

$$F_0 = \frac{F_B F_R}{F_R + F_B} \,. \tag{12}$$

Incorporating solution (11) into Eq. (10) with $v_B = \frac{dx_B}{dt}$ we recover for the bacterial sliding distance during pilus retraction

$$x_B(t) = \frac{F_0}{F_B} \left[ v^0 t + \frac{F_0}{Y} \left( e^{-\frac{Y v^0}{F_0} t} - 1 \right) \right] . \tag{13}$$

While retracting the pilus will detach with a rate constant $k_{off}(F)$ from the substrate. Assuming a force independent off-rate constant $k_{off} = k^0_{off}$ the detachment times are distributed exponentially with mean $1/k^0_{off}$. Furthermore, we assume that the single effective pilus considered in our model retracts with frequency $k_p$ and thus gives rise to an effective velocity

$$v_{eff} = k_p \langle x_B \rangle = k_p k^0_{off} \int_0^\infty x_B(t)\, e^{-k^0_{off} t}\, dt = V_{max} \frac{E}{E + E_0} \,. \tag{14}$$

Here, $\langle x_B \rangle$ denotes the mean bacterial sliding distance per pilus retraction event. $V_{max}$ denotes the maximum effective speed that a cell can reach on a given substrate at infinite rigidity, given by

$$V_{max} = v^0 \frac{k_p}{k^0_{off}} \frac{F_R}{F_B + F_R} \,. \tag{15}$$

$E_0$ denotes the rigidity at half-maximal speed and is given by

$$E_0 = \frac{F_B F_R k^0_{off}}{(F_B + F_R) v^0 \lambda} \,. \tag{16}$$

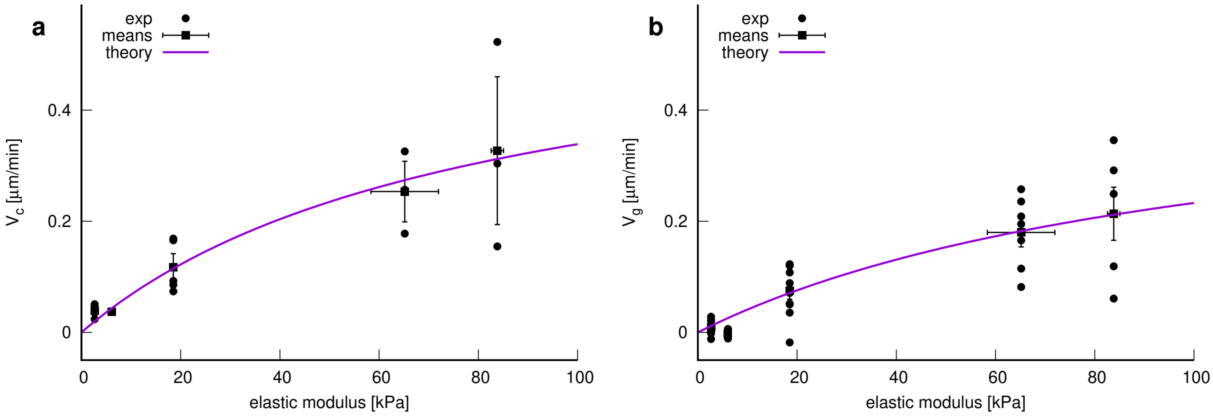


FIG. 1. Experimentally measured velocity vs. rigidity data and least squared fits of Eq. (14) w.r.t. to the experimental values as indicated in the legends. a: Local velocity measures. Parameters obtained by a least-square fit: $E_0 = 79 \pm 82$ kPa, $V_{max} = 0.61 \pm 0.35$ µm.min$^{-1}$. b: Global velocity measures. Parameters obtained by a least square fit: $E_0 = 108 \pm 101$ kPa, $V_{max} = 0.49 \pm 0.28$ µm.min$^{-1}$. Errorbars indicate SEM.

Fig. 1 shows the experimental data and fitted curves, which capture well the data for medium and high rigidities. The theoretical curves were fitted to all experimental values (applying the statistical weight in the measured rigidities and equal weight in the velocities) using a least square fit (software gnuplot [77]). Assuming a typical pilus retraction speed $v^0 = 0.5 - 1\,\mu\text{m.s}^{-1}$ [75, 76], a stall force of the order $F_\text{R} = 50 - 100\,\text{pN}$ [75, 76], a pilus off-rate constant $k^0_\text{off} = 1\,\text{s}^{-1}$ [73] and a contact size of $\lambda = 1\,\text{nm}$ [76], a high friction surface with $F_\text{B} = 1\,\text{nN}$ and a typical pilus retraction frequency [78] of $k_\text{p} = 0.1 - 0.2\,\text{s}^{-1}$ we recover a $V_\text{max} \sim 0.1 - 1\,\mu\text{m.min}^{-1}$ and a substrate rigidity at half maximum speed of $E_0 = 10 - 100\,\text{kPa}$, a range which is encloses the fitted values (see Fig. 1).

Here we have assumed a force-independent off-rate constant for the pilus. In a more complex scenario, the contact between the pilus and the substrate may act as a slip bond or catch bond. For completeness we will show some numerical results for slip and catch bond behavior below, which do not increase however the quality of fit between experimental and theoretical velocity data.

##### D. Force dependent detachment rate constants

Increasing the complexity of the model, we assume that the pilus detachment rate is force dependent [68, 73, 79, 80] and takes the form

$$k_\text{off} = k^0_\text{off}\left(\varepsilon e^{-\frac{F}{F_\text{C}}} + e^{\frac{F}{F_\text{S}}}\right). \tag{17}$$

$\varepsilon = 0$ denotes a slip bond and $\varepsilon > 0$ denotes a catch bond behavior. $F_\text{C}$ and $F_\text{S}$ denote positive force constants [68]. Eq. (17) implies that the pilus detachment times are not distributed exponentially.

We now consider the evolution equation for the probability density $p(u)$ that a pilus attached to the substrate is retracting and is thereby inducing a displacement $u$

$$\partial_t p = -k_\text{off}(F)p - \partial_u j_\text{u} \tag{18}$$

The first term denotes (tension dependent) pilus detachment from the substrate and the second term captures the advection of the displacement due to pilus retraction and bacterial sliding. It is formulated as a divergence of a flux $j_\text{u}$ with

$$j_\text{u} = [v_\text{R}(F) - v_\text{B}(F)]\,p\,. \tag{19}$$

The pilus retraction $v_R(F)$ and bacterial sliding speed $v_B(F)$ is given by Eqs. (9) and (10). To facilitate the analysis of the equations we use the transformation $p(u) = p(u[F]) = P(F)$ and $\partial_u = Y\partial_F$ which gives rise to the evolution equation

$$\partial_t P(F) = k_\text{off}(F)P - v^0 Y\partial_F\left[\left(1 - \frac{F}{F_\text{R}} - \frac{F}{F_\text{B}}\right)P\right] \tag{20}$$

To reduce the number of parameters we introduce the timescale $t_0 = 1/k^0_\text{off}$, the length scale $l_0 = v^0 t_0$ and the force scale $F_0 = (F_\text{R}F_\text{B})/(F_\text{R} + F_\text{B})$. The adimensional quantities are then denoted $\tilde{F} = F/F_0$, $\tilde{t} = t/t_0$, and $\tilde{u} = u/l_0$. The adimensional evolution equation of $\tilde{P}(\tilde{F})$ takes the form

$$\partial_t\tilde{P} = -\kappa(\tilde{F})\tilde{P} - \mu\partial_{\tilde{F}}\left[\tilde{P}(1 - \tilde{F})\right], \quad \text{with} \quad \tilde{F} \in (0, 1] \tag{21}$$

where $\mu = Yv^0/(F_0 k^0_\text{off})$ denotes the adimensional substrate rigidity and $\kappa$ denotes an adimensional force dependent off-rate, i.e. $\kappa = k_\text{off}/k^0_\text{off}$. Solving Eq. (21) in the steady state we find

$$\tilde{P} = \frac{\tilde{P}_0}{1 - \tilde{F}}\, e^{\frac{\mathcal{I}(\tilde{F})}{\mu}} \tag{22}$$

with

$$\mathcal{I}(\tilde{F}) = \varepsilon e^{-\frac{1}{\tilde{F}_\text{C}}}\text{Ei}\left(\frac{1 - \tilde{F}}{\tilde{F}_\text{C}}\right) + e^{\frac{1}{\tilde{F}_\text{S}}}\text{Ei}\left(-\frac{1 - \tilde{F}}{\tilde{F}_\text{S}}\right). \tag{23}$$

In Eq. (23) the force constants $\tilde{F}_\text{C}$ and $\tilde{F}_\text{S}$ have been rescaled by $F_0$. The normalization factor $P_0$ is defined by the integral

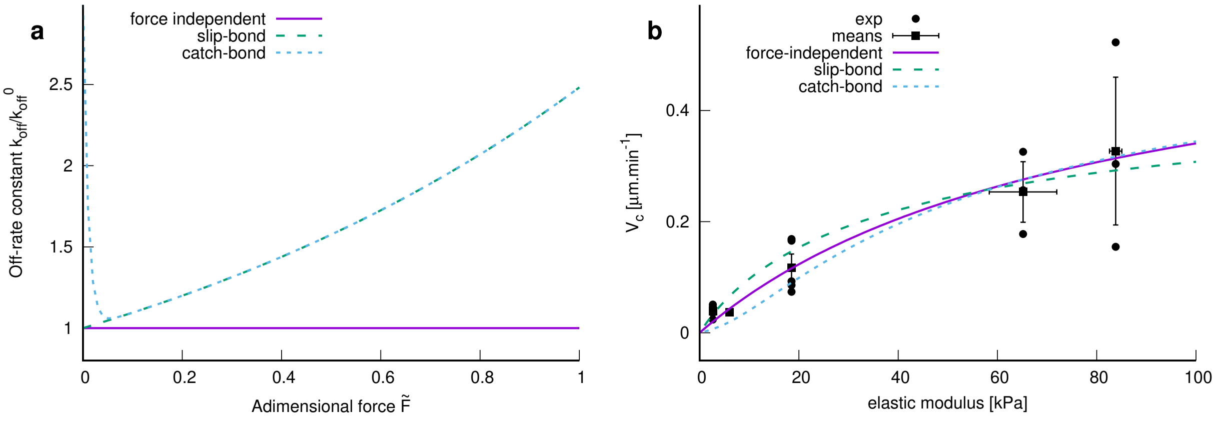


FIG. 2. a: Comparison of various force dependencies of the pilus detachment rate constant $k_{off}$ as indicated in the legend. b: Comparison of bacterial velocities obtained by models with various complexity with experimentally measured values (local velocity analysis) as indicated in the legend (parameters were fit w.r.t experimental mean values). The model parameters for the force-independent model are as in Fig. 1a. For the slip and catch bond model the fixed parameters are $E_0 = 80\,\text{kPa}$, $\tilde{F}_S = 1.1$, $\varepsilon = 2$ (catch bond), $\tilde{F}_C = 0.01$ (catch bond). For the slip and catch bond model $V_{max}$ was estimated from least square fits: $V_{max} = 1.16\,\mu\text{m.min}^{-1}$ (slip) and $V_{max} = 1.20\,\mu\text{m.min}^{-1}$ (catch). Error estimates are expected to be of the same order of magnitude as for the force independent model (see Fig. 1a).

condition $\int_0^1 \tilde{P}(\tilde{F})\,\mathrm{d}\tilde{F} = 1$. Ei$(x)$ denotes the exponential integral. At detachment the distribution of forces $\tilde{F}_d$ is given by

$$\tilde{P}_d(\tilde{F}_d) = \tilde{P}_{d0}\frac{\kappa(\tilde{F}_d)}{1-\tilde{F}_d}e^{\frac{I(\tilde{F}_d)}{\mu}} \tag{24}$$

with the normalization factor $\tilde{P}_{d0}$ determined by the integral condition $\int_0^1 \tilde{P}_d(\tilde{F}_d)\,\mathrm{d}\tilde{F}_d = 1$.

Using the forces and bacterial sliding distance at detachment from the substrate

$$\tilde{F}_d = 1 - e^{-\mu\tilde{t}_d} \tag{25}$$

$$\tilde{x}_B = \frac{1}{\tilde{F}_B}\left[\tilde{t}_d + \frac{1}{\mu}\left(e^{-\mu\tilde{t}_d} - 1\right)\right] \tag{26}$$

we can perform the transformation $\tilde{P}_d(\tilde{F}_d)\,\mathrm{d}\tilde{F}_d = \tilde{P}_d[\tilde{F}_d(\tilde{t}_d)]\mu e^{-\mu\tilde{t}_d}\,\mathrm{d}\tilde{t}_d = \tilde{P}_{\tilde{t}_d}(\tilde{t}_d)\,\mathrm{d}\tilde{t}_d$ and recover the mean bacterial displacement per pilus retraction in adimensional form as

$$\langle\tilde{x}_B\rangle = \int_0^\infty \tilde{x}_B(\tilde{t}_d)\tilde{P}_{\tilde{t}_d}\,\mathrm{d}\tilde{t}_d\,. \tag{27}$$

Following the same argument as for Eq. (14), the effective bacterial speed (dimensional) is then given by

$$v_{eff} = k_p l_0 \langle\tilde{x}_B\rangle = \frac{k_p}{k_{off}^0}v^0\langle\tilde{x}_B\rangle \tag{28}$$

$$= V_{max}\int_0^\infty \left[\tilde{t}_d + \frac{1}{\mu}\left(e^{-\mu\tilde{t}_d} - 1\right)\right]\tilde{P}(\tilde{t}_d)\,\mathrm{d}\tilde{t}_d\,, \tag{29}$$

with $\mu = E/E_0$.

Fig. 2 shows exemplarily the off-rate constants for force independent, slip and catch bond behavior (Fig. 2a) and the effective velocity of a slip-bond and catch-bond model along with a force independent detachment in comparison to the measured bacterial velocity using the local velocity analysis (Fig. 2b). Thereby we chose arbitrarily a slip-bond constant $F_S = 1.1F_0$ corresponding for example to the case of a high friction substrate with $F_R = F_S = F_B/10$, i.e. as used previously $F_R = F_S = 100\,\text{pN}$ and $F_B = 1\,\text{nN}$. The catch-bond force constant was chosen to be small, i.e. $F_S \ll F_0$, following the idea of Ref. [73] that pilus-substrate attachment is stabilized for small pilus tension. Futhermore, we chose $\varepsilon = 2$, i.e. pilus detachment at zero loads is three times faster than for a slip-bond model. Fixing $E_0 = 80\,\text{kPa}$ [obtained from fitting the force-independent model (see Fig. 1a)]., the theoretical curves with the force-dependent off-rate constant were fitted using a least square fit in the parameter $V_{max}$. The catch-bond behavior captures qualitatively better the velocities at low rigidities but neither slip-bond nor catch-bond seem to perform better than the simple analytical force-independent detachment model for medium and high rigidities.

#### III. INFLUENCE OF BACTERIAL MOTILITY ON THE ONSET OF BIOFILM VERTICALIZATION

##### A. Simple kinetic model

As described in the main text, we propose a simple kinetic model to capture the 2D to 3D transition of bacteria in growing microcolonies over time, i.e. we assume that colony verticalization results from a competition between bacterial division and motility, rather than from a competition between adhesion forces between bacteria or between bacteria and the substrate. We thus assume that there is no strong difference in the binding energy of a cell to the substrate as a function of its rigidity, and that this energy is slightly higher than that of binding to another cell. Bacteria thus prefer adhering to the substrate in all cases but can easily adhere to other cells if needed. Based on this assumption, we consider two key features of surface colonization to describe the 2D to 3D transition:

- Growth: initially, at time $t = 0$, one bacterium is attached to the surface. The number of bacteria $N$ grows exponentially with time as:

$$N(t) = e^{\frac{t}{t_0}} . \tag{30}$$

  The characteristic time scale, $t_0$, accounts both for the growth and for the occasional detachment of bacteria from the surface. *De novo* attachment of bacteria to the surface is neglected. Furthermore, $t_0$ is assumed to be constant over time and across the different surfaces.

- Movement: bacteria explore the surface with a characteristic velocity $V_{CM}$ and perform a random walk (we consider time and length scales larger than the persistence length/time of bacterial twitching motion). These displacements result in a spreading of the colony over a characteristic area $a(t)$ following a diffusive process:

$$a(t) = a_0 + \alpha V_{CM} t \tag{31}$$

  where $a_0$ is the area of one bacterium and $\alpha$ is a phenomenological parameter related to the properties of the random walk.

From the two equations above, it is clear that the number of bacteria attached to the surface grows faster than the size of the corresponding colony. Therefore, at a critical time $t_c$ corresponding to a critical number of bacteria $N_c$ on the substrate, the area available to bacteria for spreading on the surface will be completely occupied. i.e. $a_0 N_c = a_0 N(t_c) = a(t_c)$ and thus

$$N_c = 1 + \gamma V_{CM} \ln(N_c) \tag{32}$$

where we have substituted $\ln(N_c)$ for $t_c$ on the *r.h.s.* of Eq. (32) and $\gamma = \alpha\, t_0/a_0$. Solving this equation permits to obtain $N_c$ as a function of $V_{CM}$ and one unknown parameter, $\gamma$. Note that when $V_{CM} = 0$, a situation in which the bacteria do not move at all, the 2D to 3D transition occurs at the first division, i.e. as soon as $N_c > 1$.

It should be noted that here, just as $V_{CM}$ is a characteristic velocity and not the mean speed of the bacteria (see main text, Fig. 3A), that the characteristic area $a(t)$ accessible to the bacteria in the colony at time $t$ is not necessarily equal to the whole colony area: first because of their finite center-of-mass velocity $V_{CM}$; secondly, because the local density may restrict their movement and the accessible surface. This effect is difficult to quantify because the fluctuations of density inside the colony area may be, depending on $V_c$, much greater than the ones encountered in the case of the Brownian diffusion of particles. Indeed, some bacteria remain static while others explore the surface extensively (main text, Fig. 3E). Another reason is that upon division, the two daughter cells are touching and there is hence a systematic fluctuation of density upon division. Therefore the area accessible for bacteria is rather an effective measure, which cannot be directly derived from microscopic diffusion processes only. While other expressions could be used, this one is the simplest that can be proposed and matches our experimental data sufficiently well. One justification is that the underlying assumption that the velocity of bacteria is not affected by the local density (retaining the linear scaling of $a(t)$ with $V_{CM}$) is justified in the assessed situation where groups of closed-packed bacteria never exceed 5-8 cells before the 2D to 3D transition occurs. However, as a comparison, sub- and super-linear scalings will be compared with the data in Subsection III B.

To further analyze the microscopic meaning of $\gamma$, we note that it is the inverse of a velocity and is related to the compactness of the colony, with higher values indicating a sparser distribution of bacteria with a lesser probability that growing/twitching bacteria will encounter several others and move to 3D because of local crowding. However it is misleading to compare it to values that could be derived from random walks with persistence because of the above-mentioned discrepancy between the colony area and the area accessible to bacteria for further spreading. Relating $\gamma$ to experimentally measured quantities on the cell movement would require a detailed analysis of the cell density fluctuations on the surface which is beyond the scope of this paper.

##### B. Comparison of experimental data with the model

All available data from which characteristic velocities were extracted (main text, Fig. 3B) were analysed and included, with the exception of one data point on glass due to the presence of an air bubble on the surface before the onset of the 2D→3D transition. The characteristic velocities for each experiment and each rigidity were taken from Fig. 3B. The characteristic number of bacteria $N_c$ per colony was estimated as follows.

- First, for low- and medium-ridigity surfaces (2.7 kPa, 6.1 kPa and 18.5 kPa), colony formation from isolated bacteria was monitored over time until the 2D to 3D transition occurs. The number of bacteria on the surface stemming from the initial isolated bacteria were then counted, and the count for all the colonies were averaged to calculate $N_c$. In addition, the average number of colonies forming in the observed area up to that point was also measured.

- For higher-rigidity surfaces, the movement of bacteria is too large to keep track of all bacteria stemming from the same progenitor as they mix or leave the field of view, while others are incoming. As a result, $N_c$ was calculated by counting the total number of bacteria in the field of view at the time of the onset of the 2D to 3D transition, and dividing this number by the estimated number of colonies as measured on low-rigidity surfaces. It should be noted that in this case, the simple model presented above is not valid as it considers only one isolated colony, and can be expected to yield overestimated values of $N_c$. Furthermore, our evaluation method of the number of colonies in the field of view may be prone to error so we used a "blind" evaluation procedure performed before the count of bacteria in the field of view, to avoid possible biases. A change of 1 (compared to a mean value around 4) in the number of colonies used to normalise the total number of bacteria provides a good estimate of the error bars on each individual data point, and is comparable to the spread of the data points (see Fig.3). When several surfaces of the same rigidity have been measured in one experiment, the different $N_c$ values are averaged.

The above cited procedure produced one doublet ($V_{CM}$, $N_c$) for each rigidity of each experiment.

To match our simple model with the experimental data, Eq. (32) can be used to calculate $N_c$ as a function of $V_{CM}$ for a given value of $\gamma$. However, a direct fit of the experimental data is difficult as there is no analytical solution to Eq. (32). Instead, an experimental value of $\gamma$ was calculated from each experimental point using the expression

$$\gamma_{exp} = \frac{N_c - 1}{V_{CM} \ln(N_c)} \tag{33}$$

An average experimental value is then calculated, along with a standard deviation, $\langle\gamma_{exp}\rangle = 56.8$ min.µm$^{-1}$ and $\delta\gamma = 11.2$ min.µm$^{-1}$.

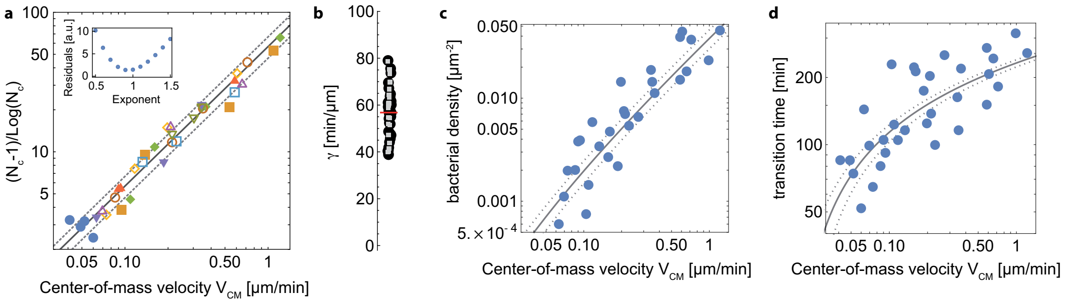


FIG. 3. *a, comparison between the experimental data (markers) and the kinetic model (lines). Blue dots are data obtained with the pili-deficient mutant pilA : Tn5. Inset, residual of the fit of all the experimental data points as a function of the exponent value for $V_{CM}$, indicating that the best fit is achieved for a value of or close to 1. b, $\gamma_{exp}$ values for all data points (black disks) and their average (red line). Gray squares are data points with $v_0 < 2\,\mu m.min^{-1}$, showing a similar distribution and thus ruling out a significant bias at high velocities. c, same dataset as in a but the surface density at the transition is plotted; d, same dataset as in a but the time at the transition is plotted. The same model is used to describe the data, but converted into the proper quantities.*

Fig. 3a shows the experimental data $(N_c-1)/Log(N_c)$ as a function of $V_{CM}$ (each marker corresponds to a different experiment), and the corresponding theoretical straight lines with slopes $\langle\gamma_{exp}\rangle$ (solid line), and $\langle\gamma_{exp}\rangle$ +/- 1 standard deviation (dotted lines). To assess the deviation from the curve at high velocities, $\langle\gamma_{exp}\rangle$ was also calculated from all data points with $v_0 < 2$ µm.min$^{-1}$ but the change in the value is minimal (58.2 min.µm$^{-1}$ instead of 56.8 min.µm$^{-1}$, see Fig. 3b).

Our strongest assumption in this modelling is the expression of $a(t)$ as a function of $V_{CM}$ [Eq. (31)]: an obvious *a posteriori* evidence for its correctness is that the derived equation fits our data well over more than one decade in velocity. To strengthen our point, however, we have also calculated similar curves using an exponent for $V_{CM}$ ranging from 0.5 to 1.5 (steps of 0.1, Fig. 3a, inset): the comparison with experimental data indicates that reasonable agreement is only obtained for exponent values between 0.8 and 1.1, at most.

Finally we would like to point out that the data do not collapse as well when plotting the density of bacteria, or the time of the 2D→3D transition (Fig. 3c and d). A likely explanation is that the initial number of bacteria on the surface varies between different datasets, a bias that is cancelled when plotting the number of bacteria instead of the density or the time at the onset of the transition. The same model is used with the same average parameter and spread, but converted into the proper quantities: for the density, the curves in Fig. 3a are multiplied by the average number of colonies per observed area (3 colonies), and divided by the image area (26121 $\mu m^2$); for the transition time, the logarithm of the number of bacteria per colony at the transition is multiplied by the typical growth time of the number of bacteria on the surface (∼40 min). This time incorporates both the division time (∼30 min) and the departure of a fraction of the bacteria from the surface.

#### IV. SUPPLEMENTARY FIGURES

##### A. Behavior of pili-deficient mutant

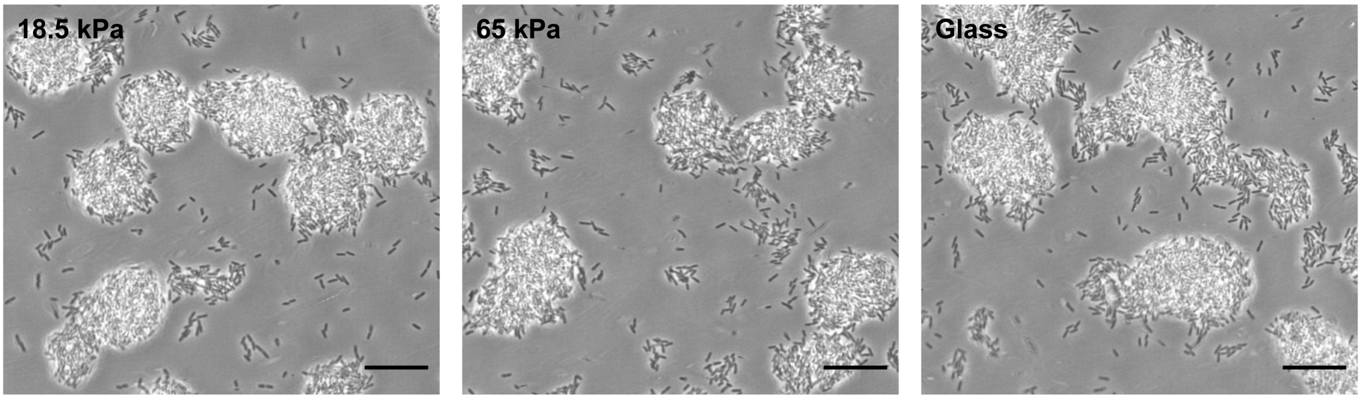


FIG. 4. In the T4P-deficient mutant $PAO1pilA : Tn5$, substrate rigidity does not significantly impact colony morphology. Colonies imaged after 10 h. Scale bar 20 µm.

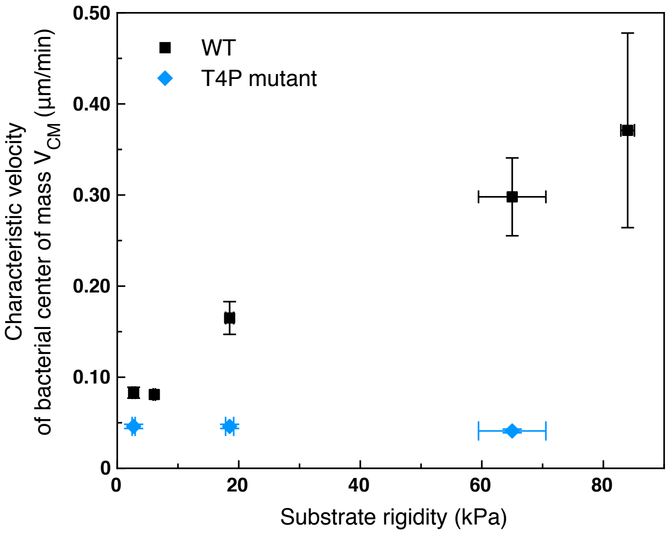


FIG. 5. Characteristic velocities of the bacterial center of mass $V_{CM}$ measured via full-tracking analysis, for the WT PAO1 strain and a T4P-deficient mutant.

#### B. Characteristic velocity on PAA and PEG gels

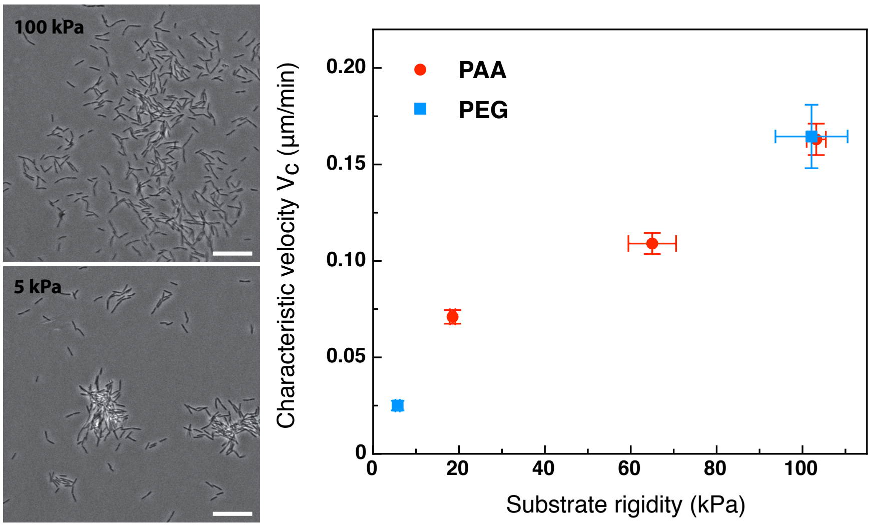


FIG. 6. Characteristic twitching velocity measured on PEG hydrogels is similar to the one measured on PAA hydrogels under identical experimental conditions. Left, phase contrast images of wt PAO1 bacteria on PEG hydrogels 5h after the onset of surface colonization. As for PAA hydrogels, the morphology of microcolonies is strongly impacted by surface rigidity. Scale bars, 20 µm. Right, characteristic twitching velocity $V_c$ ($V_{CM}$ obtained by fitting the center of mass distribution, minus $V_{CM}$ measured for the T4P-deficient mutant) on PAA or PEG hydrogels in a similar range of substrate rigidity. Error bars are SEM. Each point is the average of 2 positions from 1 experiment.

##### C. Distributions of path durations and mean track velocities

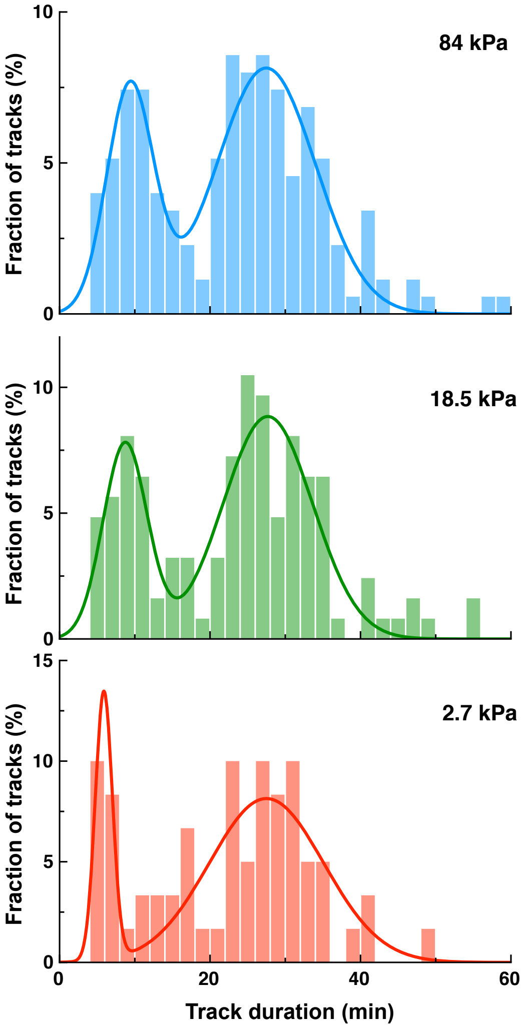


FIG. 7. Distribution of the path duration on different substrates for WT PAO1, and bimodal gaussian fit.

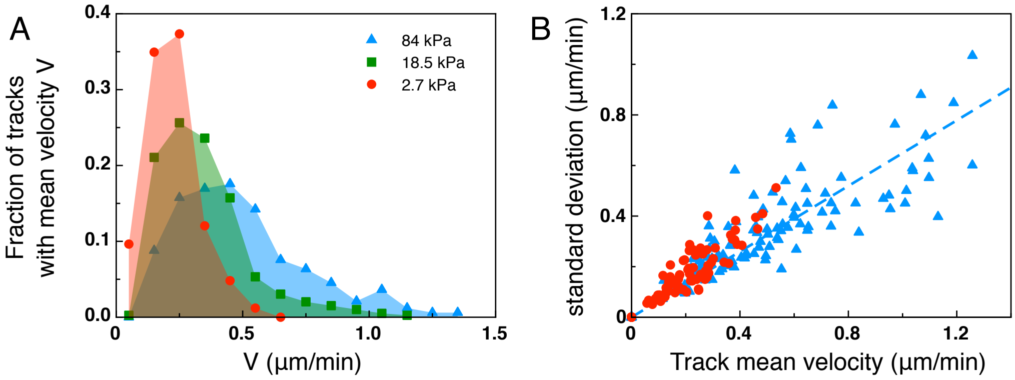


FIG. 8. Anaysis of the mean track velocity. (A) Mean track velocity distribution for different values of the substrate rigidity. Only full tracks were considered (corresponding to the right peak in Fig.7). Considering all tracks does not significantly modify the distributions (data not shown). 84 kPa: 330 tracks from 2 independent experiments, 18.5 kPa: 394 tracks from 3 independent experiments, 2.7 kPa: 83 tracks from 2 independent experiments. (B) Standard deviation of the mean velocity for individual tracks. Dotted line is a linear fit of the data for 84 kPa (y=0.65x).

##### D. Extracellular matrix deposits

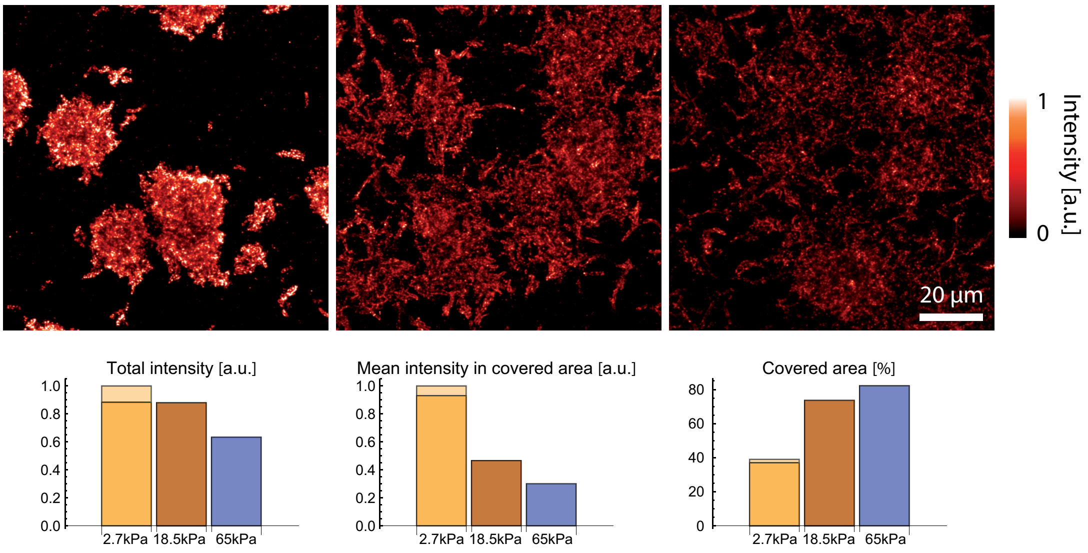


FIG. 9. EPS staining with concanavalin A highlights matrix deposits on the surface of substrates of different stiffnesses after 8h of surface colonization. Matrix deposits are more compact on soft substrates with large areas (>60%) devoid of EPS, while stiffer substrates are almost fully decorated (>% covered area). The two colours for the softer surface corresponds to two areas taken 30 minutes apart after staining, before and after acquisition of the two other datasets on larger rigidities, to rule out any significant time evolution of the staining.

##### E. Mixing of strains during surface co-colonization

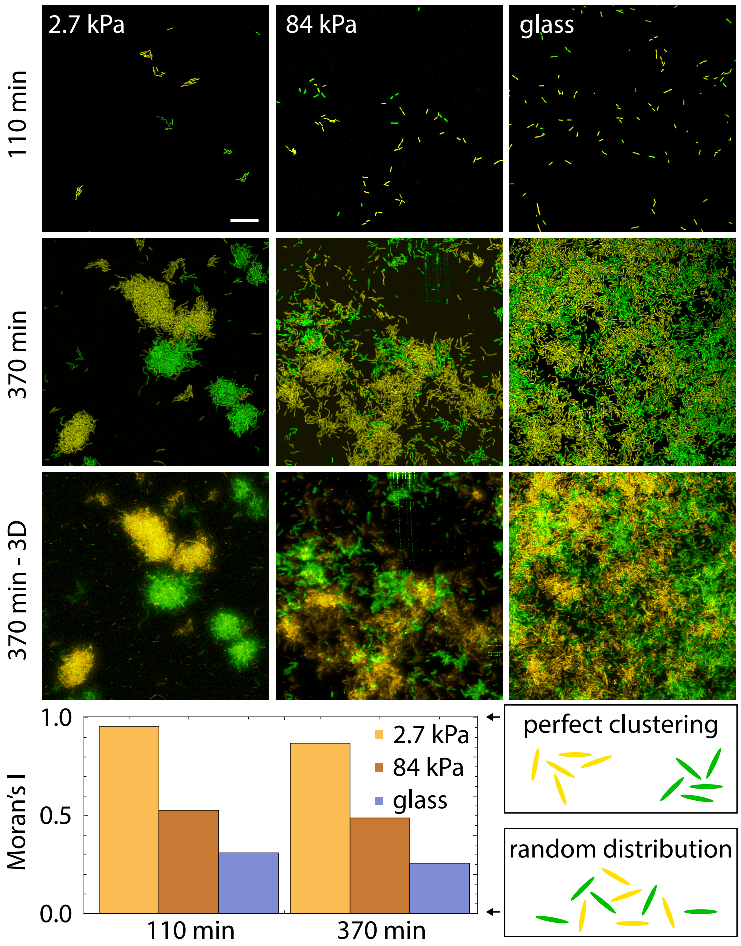


FIG. 10. Bacterial spatial distribution as a function of substrate rigidity. Top, images of surfaces seeded with a 1:1 mixture of constitutively fluorescent bacteria expressing GFP or YFP at two different times after the start of surface colonization (bottom : 3D rendering of a volumetric image). Bottom, Spatial autocorrelation quantified via Moran's I index at the two time points illustrated above. The values should not be directly compared between the two time points (see SI I-C) but illustrate that the rigidity modulation of mixing is maintained over time from very sparse to large coverage of the surface.

#### F. Time evolution of crdA expression

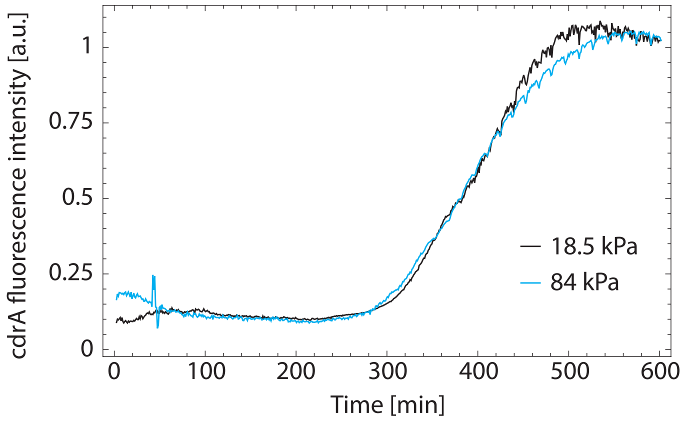


FIG. 11. fluorescence intensity from a PAO1 strain expressing a $P_{cdrA} - gfp$ plasmid over time, showing a plateau of fluorescence expression after ≈500 min. In our experiments, this change in behaviour might be due to a lack of oxygen in the flow upon growth of the biofilm in the microfluidic channel and occurs. Two rigidities are shown and scaled to the same final fluorescence value.

#### G. Mechanical characterization of the gels

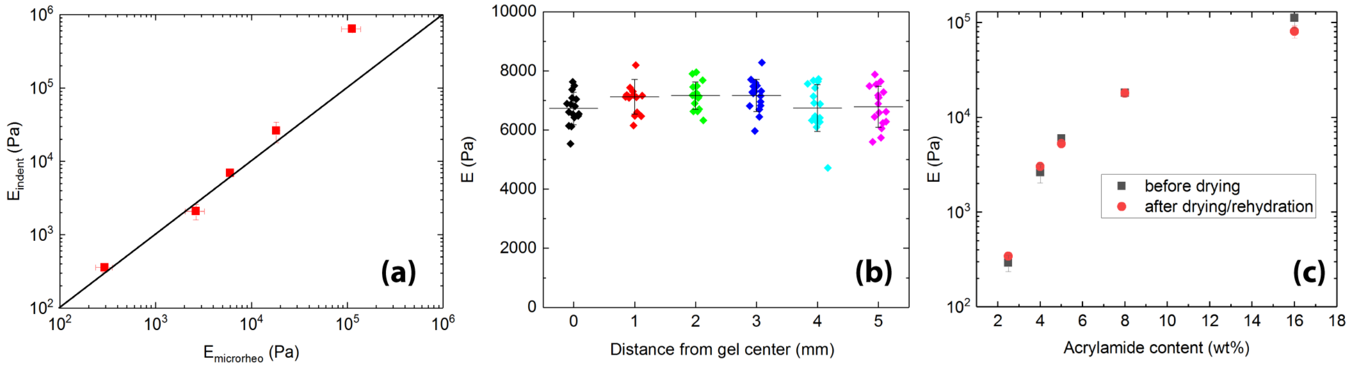


FIG. 12. (a) comparison of elastic moduli measured by indentation and by microrheology. Both techniques are seen to yield quantitatively similar results for gels with Young's moduli $E < 20$ kPa. (b) spatial homogeneity of the gels characterized by indentation measurements. For each position, separated by 1 mm, a 4x4 force spectroscopy map is taken, with a spacing between "pixels" of 3 µm. (c) comparison of elastic moduli $E$ measured before and after drying of the gels.

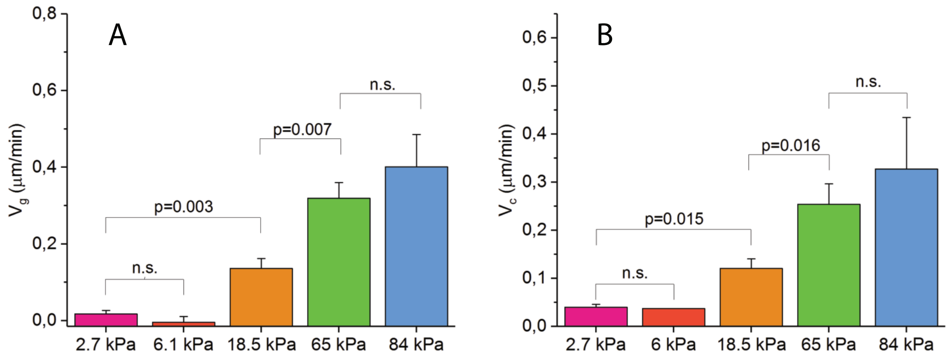


FIG. 13. Statistical analysis of $V_g$ (A) and $V_c$ (B) for various gel rigidities. Indicated are p-values from paired two-sided Mann-Whitney non-parametric tests. n.s. denotes datasets that are not significantly different at a threshold of 0.05.

[59] P. Thevenaz, U. E. Ruttimann, and M. Unser, IEEE transactions on image processing **7**, 27 (1998).
[60] I. Arganda-Carreras, V. Kaynig, C. Rueden, K. W. Eliceiri, J. Schindelin, A. Cardona, and H. Sebastian Seung, Bioinformatics **33**, 2424 (2017).
[61] J.-Y. Tinevez, N. Perry, J. Schindelin, G. M. Hoopes, G. D. Reynolds, E. Laplantine, S. Y. Bednarek, S. L. Shorte, and K. W. Eliceiri, Methods **115**, 80 (2017).
[62] P. A. Moran, Biometrika **37**, 17 (1950).
[63] L. D. Landau and E. M. Lifschitz, Theory of Elasticity, 3rd ed., Theoretical Physics, Vol. 7 (Butterworth Heinemann, 2004).
[64] A. N. Simsek, A. Braeutigam, M. D. Koch, J. W. Shaevitz, Y. Huang, G. Gompper, and B. Sabass, Soft Matter **15**, 6224 (2019).
[65] P. Sens, EPL (Europhysics Letters) **104**, 38003 (2013).
[66] P. Robert, A.-M. Benoliel, A. Pierres, and P. Bongrand, Journal of Molecular Recognition: An Interdisciplinary Journal **20**, 432 (2007).
[67] E. Evans, Annual review of biophysics and biomolecular structure **30**, 105 (2001).
[68] Y. V. Pereverzev, O. V. Prezhdo, M. Forero, E. V. Sokurenko, and W. E. Thomas, Biophysical Journal **89**, 1446 (2005).
[69] J. Kuik, S. Vincent, B. Leeflang, L. Kroon-Batenburg, and J. Kamerling, Canadian Journal of Chemistry **84**, 730 (2011).
[70] J. F. Marko and E. D. Siggia, Physical Review E **52**, 2912 (1995).
[71] J. M. Skerker and H. C. Berg, Proc. Natl. Acad. Sci. USA **98**, 6901 (2001).
[72] M. D. Koch, C. Fei, N. S. Wingreen, J. W. Shaevitz, and Z. Gitai, Proceedings of the National Academy of Sciences **118**, e2014926118 (2021).
[73] L. Talà, A. Fineberg, P. Kukura, and A. Persat, Nat. Microbiol. **4**, 774 (2019).
[74] A. Beaussart, A. E. Baker, S. L. Kuchma, S. El-Kirat-Chatel, G. A. O'Toole, and Y. F. Dufrêne, ACS nano **8**, 10723 (2014).
[75] R. Marathe, C. Meel, N. C. Schmidt, L. Dewenter, R. Kurre, L. Greune, M. A. Schmidt, M. J. Müller, R. Lipowsky, B. Maier, and S. Klumpp, Nature Communications **5** (2014), 10.1038/ncomms4759.
[76] M. D. Koch, E. Han, J. W. Shaevitz, and Z. Gitai, bioRxiv (2021), 10.1101/2021.08.26.457786, https://www.biorxiv.org/content/early/2021/08/26/2021.08.26.457786.full.pdf.
[77] T. Williams, C. Kelley, and many others, "Gnuplot 5.2: an interactive plotting program," `http://gnuplot.sourceforge.net/` (2019).
[78] Here we assume that one single effective pilus is active during a retraction event. Using a typical pilus length of 5 μm with retraction speed of $v_0 = 0.5 - 1$ μm.s$^{-1}$ gives a duration of 5-10 s per retraction and a retraction frequency of 0.1-0.2 s$^{-1}$.
[79] H. Kramers, Physica **7**, 284 (1940).
[80] O. Björnham and O. Axner, Biophysical Journal **99**, 1331 (2010).
